# Unveiling the Toxicological and Allelopathic Effects of *Pteridium aquilinum*: Chemical Profiling and Biological Assays

**DOI:** 10.64898/2026.01.09.698682

**Authors:** Isabel Feito, Alexis E. Peña, L. María Sierra, José Manuel Alvarez, Mª Lucía Rodríguez, Bernardo Barredo, Iván Iglesias, Ana Velázquez, Andrea Bartolomé, Daniel Díaz, Ruth K. Hernández, Juan Majada, Elena Canga, Helena Fernández

## Abstract

The fern *Pteridium aquilinum* (bracken) is among the world most widespread plants and poses serious ecological and health risks due to the presence of illudane glycosides (IGs), carcinogenic compounds that particularly affect grazing livestock. This study determined the presence of IGs in plant extracts and in water samples from the Asturias region (northern Spain), an area characterized by intensive cattle farming, and assessed the genotoxic potential, and allelopathic effects of these samples. The glycosides ptaquiloside (PTA) and ptesculentoside (PTE), together with the degradation products pterosin A and pterosin B, were quantified in *in vitro*-cultured gametophytes and young sporophytes, spring-collected croziers, and autumn water samples from several bracken-infested sites. All sampling locations contained IGs in both plant and water samples, with particularly high concentrations of PTA detected in croziers from Brana Vieja (Somiedo), where cattle deaths had been reported. *In vivo* genotoxicity assays, using *Drosophila melanogaster*, revealed induction of somatic mutations and recombination events by aqueous plant extracts and of somatic mutations by water samples. The genotoxic activity of plant extracts, but not that of water samples, was associated with pterosins A and B. Moreover, comet assays in cultured human cells confirmed the genotoxic activity of plant extracts. Unexpectedly, allelopathic bioassays indicated a possible phytostimulatory rather than inhibitory effect of bracken extracts on the germination of meadow species. These findings underscore the widespread presence and biological activity of bracken in northern Spain and highlight the urgent need for management strategies to mitigate the ecological and toxicological threats posed by bracken proliferation.

## 1. Introduction

*Pteridium aquilinum (L.) Kuhn,* or bracken, is one of the most widely distributed vascular plants, present on all continents except Antarctica (Marrs et al., 2010). It grows optimally at 15-30 °C in soils with pH 5.5-7.5 (Marrs & Watt, 2006). In temperate and boreal Europe, fronds emerge in spring and die back in autumn, while in tropical regions they may persist over a year (Den Ouden, 2000). Its extensive distribution is due to high spore production and vegetative reproduction, enabling rapid population expansion (Rasmussen et al., 2015).

Bracken outcompetes other plants through dense, long-lived fronds that form thick canopies and deplete soil nutrients and water (Marrs & Watt, 2006). Allelopathic effects have been observed, such as inhibition of tomato germination by bracken extracts, though not consistently (Loresco et al., 2006; Bracho & Arnaud, 2012). Control measures, including repeated mowing or herbicide application (e.g., Asulam), reduce rhizome carbohydrate reserves and limit spread (Pakeman et al., 2002; Gil da Costa et al., 2024).

Beyond biodiversity impacts, bracken is toxic. In upland and temperate grazing areas, it can form a large part of the vegetation. Livestock usually avoid it but may ingest it during food scarcity or accidentally in hay (Marrs & Watt, 2006; Rai et al., 2017). Prolonged exposure can cause severe health problems or death, depending on dose and species (Alonso-Amelot & Avendaño, 2002; Evans et al., 2009; Francesco et al., 2011; Suazo-Ortuño et al., 2015). Common syndromes include thiamine deficiency in horses, acute hemorrhagic disease in sheep and cattle, bright blindness in sheep, bovine enzootic hematuria, and carcinoma of the upper digestive tract in cows (Vetter, 2009; Gil da Costa et al., 2012; 2024)

In humans, consumption of bracken represents a significant health risk. Although rare today, ingestion of rhizomes and young fronds (croziers) still occurs in regions such as Japan, the United States, Canada, Brazil, Costa Rica, Angola, and South Africa (Ugochukwu, 2019; Malik et al., 2023). Boiling the plant reduces, but does not eliminate, its toxicity (Vetter, 2009). Regular consumers have a higher risk of cancers of the upper digestive tract: studies in Venezuela reported a 2- to 4-fold increase in gastric cancer (Alonso-Amelot & Avendaño, 2001), and Brazilian populations also show elevated incidences of esophageal and gastric cancers (Marlière et al., 2019). Humans may also be exposed indirectly through dairy and meat products from contaminated animals (Alonso-Amelot et al., 1993, 1996; Fletcher et al., 2012; Virgilio et al., 2015), inhalation of spores (Povey et al., 1996; Siman et al., 2000; Rasmussen et al., 2013; Kiselius et al., 2024) or contact with contaminated soil and water (Rasmussen et al., 2003, 2005; Schmidt et al., 2005; Jensen, 2008; Owesen et al., 2008; Clauson-Kaas et al., 2014, 2016; O’Driscoll et al., 2016; García-Jorgensen et al., 2021; Skrbic et al., 2021).

Bracken toxicity is caused by various secondary metabolites, including dihydrofolic, benzoic, cinnamic, and p-coumaric acids, thiaminases, cyanogens, glucopyranosides, braxins, flavonoids, tannins, p-hydroxystyrene derivatives, and illudane glycosides (IGs) (Vetter, 2009; O’Connor et al., 2019; Marques et al., 2024). Key IGs include ptaquiloside (PTA), caudatoside (CAU), ptesculentoside (PTE), and related compounds such as isoptaquiloside (IPTA) and ptaquiloside Z (PTA-Z) (Gil da Costa et al., 2012; O’Connor et al., 2019; Marques et al., 2024). These IGs can hydrolyze to form pterosins: PTA to pterosin B, CAU to pterosin A, and PTE to pterosin G (Gil da Costa et al., 2012; Vetter, 2009; Aranha et al., 2019; O’Connor et al., 2019). The reactive dienone intermediates generated during hydrolysis are capable of covalently binding to DNA, forming alkylated adducts that are responsible for the carcinogenic properties observed in exposed animals (Kushida et al., 1994; Povey et., 1996; Freitas et al., 2001; Alonso-Amelot & Avendaño, 2002; O’Connor et al., 2019). (**Figure 1**).

**Figure 1.**
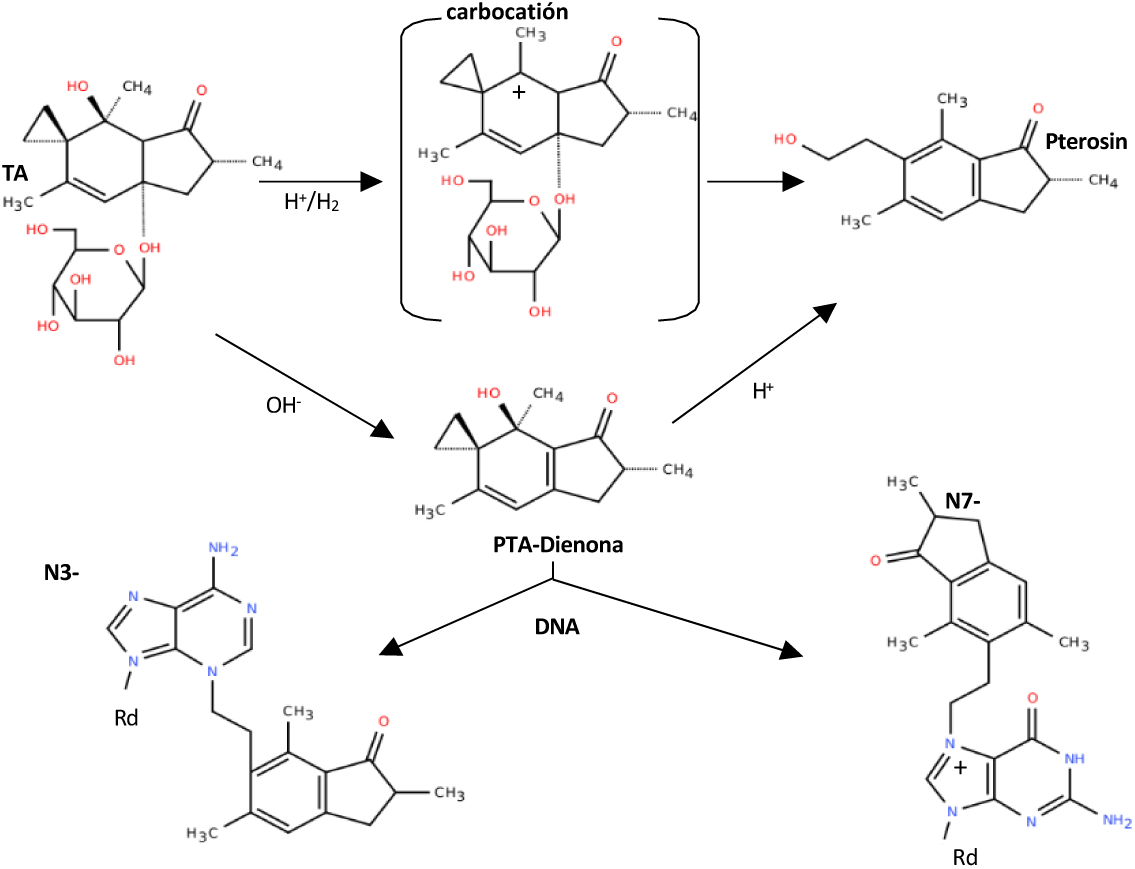
Activation and stabilization of ptaquiloside (PTA) and induced DNA adducts. Schema adapted from O’Connor, et al., (2019).

Although PTA was long considered the main carcinogen in bracken due to its abundance, recent studies have shown significant levels of CAU and PTE in *P. aquilinum* from northern Europe and *P. esculentum* from Australia (Kisielius et al., 2022). Global sampling -excluding Antarctica- indicated approximate distributions of 41% PTA, 37% CAU, and 22% PTE among Pteridium species. PTA predominates in *P. aquilinum*, whereas PTE is dominant in Australian *P. esculentum*, representing 40-73% of total illudane glycosides (Kisielius et al., 2022).

Research on bracken in Spain, and particularly in the Asturias region, remains very limited despite the ecological and health risks posed by this widespread fern (Rodriguez de la Cruz et al., 2009; García-Arroyo et al., 2015, 2017). Recently, Sierra et al. (2025) published a comprehensive multidisciplinary study, assessing toxic compounds, genotoxic activity of aqueous plant extracts, bracken risk to cattle and its distribution in northern Spain. Nevertheless, there is a clear research gap in integrating developmental stage–specific analyses, environmental contamination, and allelopathic effects in high-livestock-density regions such as Asturias, where the widespread presence of bracken fern poses a sustained and regionally relevant threat to grazing cattle. The present work extends and complements Sierra et al. (2025) to provide a more comprehensive assessment of the occurrence of illudane glycosides (IGs), and of possible biological effects of bracken across different developmental stages and environmental matrices. Specifically, we focus on *in vitro* cultured gametophytes and young sporophytes, as well as field-collected croziers in spring and fall, and autumnal water samples, to evaluate ptaquiloside, ptesculentoside, and the degradation products pterosin A and B. In addition, we investigate the potential genotoxicity, and allelopathic effects on the germination of meadow species, of bracken aqueous extracts. By integrating these multiple approaches, this study seeks to fill critical knowledge gaps regarding the ecological and toxicological impact of bracken in a region of intensive cattle farming, thereby informing rural medium responsible entities.

## 2. Material and methods

### 2.1 *In vitro* culture of spores of *P. aquilinum*

Fronds of *P. aquilinum* with sporangia were collected from Las Golondrinas Trail (14°10ʹ41.86ʺ N; 87°03ʹ21.15ʺ W), located in the municipality of Valle de Ángeles, La Tigra National Park, Honduras. The fronds were air-dried at ambient temperature, ground using an electric grinder, and the resulting powder was stored at room temperature until use. Prior to *vitro* culture, 0.5 mg of this powder were placed in 15 mL PYREX centrifuge tubes, hydrated for 24 h, and disinfected with a commercial bleach solution (5 g/L), containing 0.1% Tween 20, for 10 min, under sterile conditions in a laminar flow cabinet. Spores were then washed three times by centrifugation at 2,500 rpm for 3 min to remove the bleach solution. Finally, the spores were resuspended in 5 mL of sterile distilled water, and 2.5 mL were transferred, using a Pasteur pipette, into 500 mL flasks containing MS liquid medium (Murashige and Skoog, 1962) supplemented with 20 g/L sucrose, adjusted to pH 5.8. The flasks were placed on an orbital shaker at 75 rpm and maintained at 25 ± 2 °C, under a 16-hour photoperiod. Three months later, germinated gametophytes were transferred to full- or half-strength MS solid medium containing 8 g/L agar in 125 mL flasks. One month afterward, data on sporophyte development, including number of fronds per flask and frond length (in cm), were recorded.

### 2.2 Obtention of illudan glycosides from Pteridium esculentum for analytical purposes

The illudane glycosides ptaquiloside (PTA), ptesculentoside (PTE), and caudatoside (CAU) were isolated and purified from *P. esculentum* plants growing in Australia (**Figure S1**). The method described by Kisielius et al. (2020) was adapted to specific laboratory conditions. For this purpose, 10 g of freeze-dried aerial plant parts were placed in a 500 mL flask with 200 mL of Milli-Q water, protected from light with aluminum foil, were subjected to ultrasound at room temperature for 30 min, and then were centrifuged for 30 min at 3500 rpm and 4 °C. The supernatant was vacuum-filtered using Whatman 114 filters, and the resulting extract was stored at –20 °C for 24 h. After thawing, the extract was centrifuged for 10 min at 3,500 rpm and 4 °C and, subsequently, filtered to remove impurities. The extract was then analyzed by HPLC-PAD (Waters 2695 with PAD 2996 detector), at a flow rate of 1 mL/min under the following conditions: injection volume 25 μL, temperature 20 °C, mobile phase MeOH:Milli-Q H₂O (55:45, pterosin method), isocratic, running time 25 min, column Kromasil 100-5 C18 (5 μm, 25 × 0.46 cm), and detection wavelengths 220 and 264 nm.

The extract was divided into two fractions of approximately 100 mL each and applied to two columns containing CC6 polyamide resin, where it was eluted at a flow rate of ∼1 mL/min. The fractions were analyzed by HPLC-PAD at 220 and 264 nm. Subsequently, the polyamide column extracts were passed through SPE-HLB cartridges, previously conditioned with 12 mL of Milli-Q water. Each cartridge received 50 mL of extract, followed by a wash with 5 mL of Milli-Q water and sequential elution with 5 mL of 40% methanol in water. The resulting eluates were analyzed by HPLC-PAD under the same conditions. Hydrolysis was performed on a portion of the polyamide column extract to identify signals corresponding to illudane glycosides. For this, 1 mL of the solution was passed through an SPE-PA cartridge, 70 μL of 1 M NaOH was added, and the mixture was vortexed and incubated in a water bath at 35 °C for 15 min. Then, 35 μL of chilled 2.5 M trifluoroacetic was added. Samples were stored at –20 °C until further analysis by HPLC-PAD. Pterosins were identified using PtrA (CAS 35910-16-8) and PtrB (CAS 34175-96-7) standards obtained from Chem Faces.

The 40% methanol–water eluate was fractionated by semi-preparative HPLC-DAD (Agilent 1260 Infinity) using a PR-C18 column under isocratic conditions (MeOH:H₂O 42.5:57.5, 3 mL/min, 27 min). Fractions were collected every 0.5 min, pooled based on chromatographic profiles, and dried by SpeedVac and freeze-drying. The isolated solids were characterized by ¹H NMR (Bruker 600 MHz) and mass spectrometry (ESI-qTOF and APCI). Compound purity was determined by quantitative ¹H NMR using an internal standard, following Kisielius et al. (2020).

### 2.3. Quantification of metabolites in croziers and water sources

The compounds PTA, PTE, PtrA, and PtrB were analyzed in plant aqueous extracts and water source samples collected from several locations in Asturias (see Table 1 and Figure 3). Specifically, plant samples (crozier tissues) were collected in the field from three areas in Asturias: the Urbiés Mountains (MU2), and the natural park of Somiedo (BRV) and the national park Picos de Europa (PEBEL). In addition, the contents of these compounds were assessed in spore-derived gametophytes and sporophytes grown *in vitro*. Regarding the water source samples, they were collected from ten locations across Asturias, from north to south and east to west, including sites within the natural parks of Fuentes del Narcea, Ibias y Degaña (DEG), Redes (RED), La Mesa-Ubiñas (LaM), Ponga (PON), and the national park of Picos de Europa (PEVF, PEBEL, PELH), as well as from the Urbiés Mountains (MU2 and MU11) and the coastal area of Pría (PRI) (see **Table 1**).

**Table 1.**
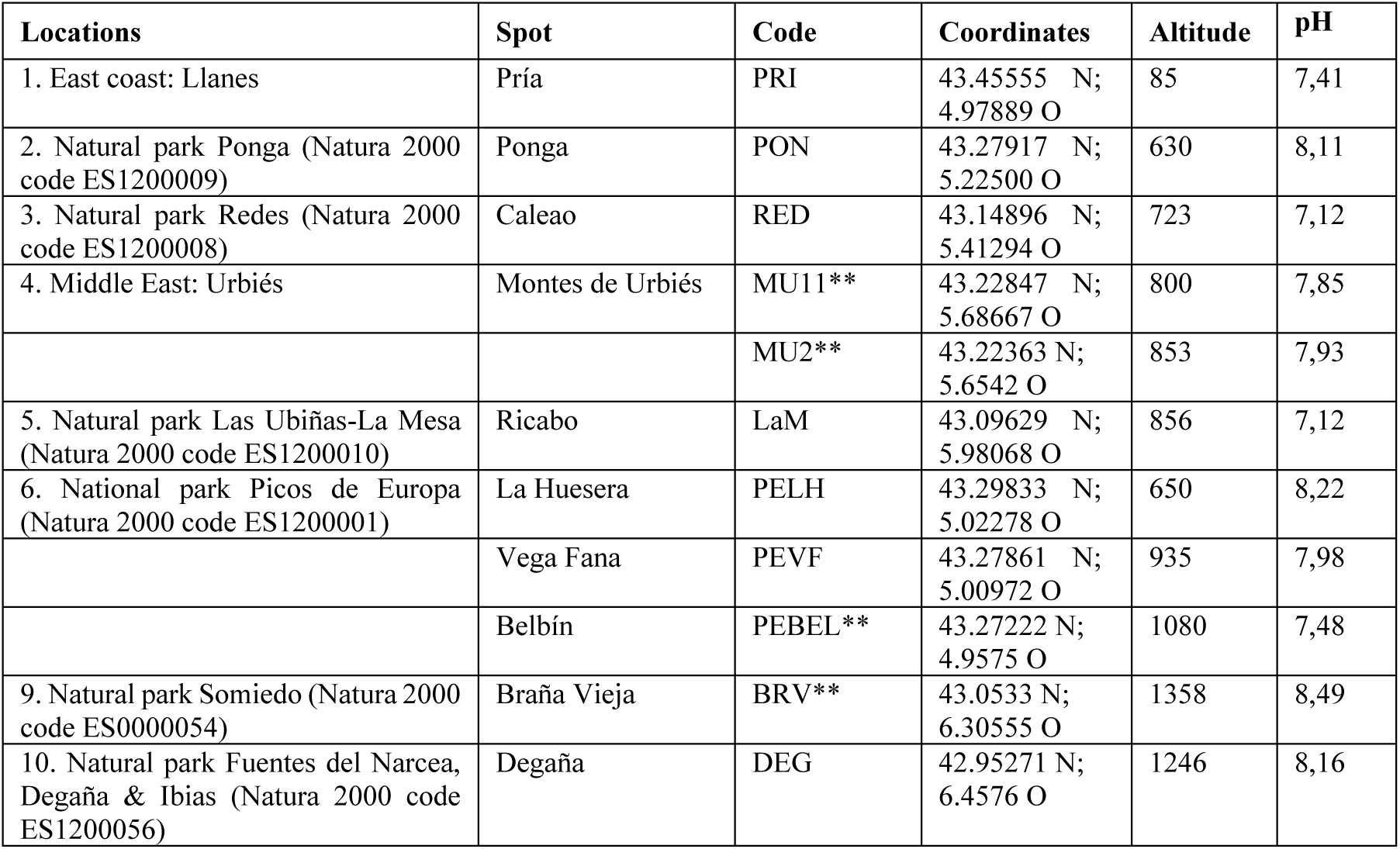
Locations and sites, and their names, for the sampling sites where water sources and croziers (in this case noted by asterisk) were collected, ordered by altitude. Physical parameters of altitude (in meters) and coordinates are included as well as the Natura 2000 codes for the national and Natural Parks, and the pH of the water samples.

**Figure 3.**
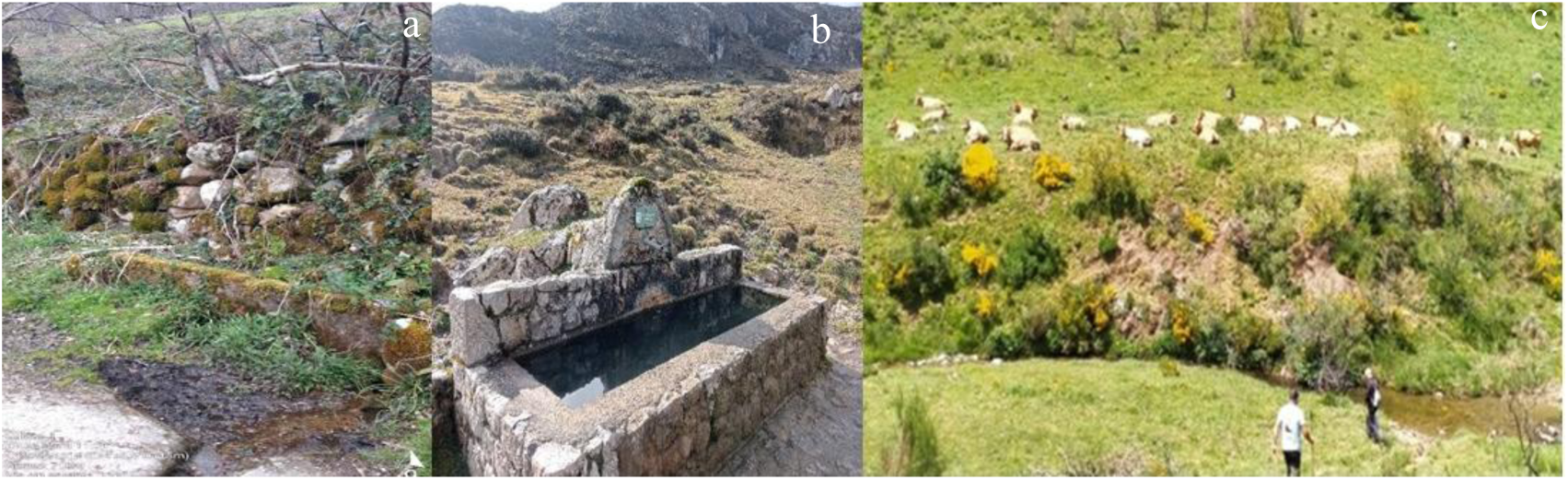
Collecting sites in areas exposed to bracken of: a, b) water sources in the Natural Park of Redes (Caleao), and the National Park of Picos de Europa (Belbín); and c) croziers in the Natural park of Somiedo, close to a cow’s resting area in Braña Vieja.

The analysis followed the protocol described by Sierra et al. (2025). Plant powder (40 mg) was extracted with 40 mL of 20% MeOH in 0.1 M ammonium acetate (pH 6), shaken at 4 °C for 20 min and centrifuged (3000 × g, 5 min). Loganin was added as an internal standard, and extracts were filtered (0.22 µm PVDF). Compounds were separated and quantified by UHPLC–MS/MS using an Agilent 6460 Triple Quad system. Separation was performed on a Poroshell 120 EC-C18 column using 0.1% formic acid in water (A) and acetonitrile (B) at 1 mL min⁻¹ under a gradient elution. Positive ESI and dMRM mode were used for detection. Three independent extracts per sample were analyzed in triplicate, with blanks between runs.

Regarding the water sources, two samples of 42,5 mL from each spot were collected, lyophilized, and then resuspended in 200 μL of initial HPLC solvent, filtrated and directly injected, following the mass analyses conditions aforementioned for croziers.

### 2.5. Genotoxicity Analysis: SMART Assay

The eye *w/w+* SMART assay was performed with the Oregon K strains *yellow* and *white* (Ok-*y* and Ok-*w*, respectively) of *D. melanogaster* and surface treatments, as described by Sierra et al. (2025). Briefly, second-third instar larvae, from mass crosses between Ok-*y* females and Ok-*w* males, and developed in Caroline Instant Drosophila medium (Formula 4-24), were treated with 1.5 mL/bottle of different solutions from the analysed samples, and the eyes of the hatched adults were scored as described (Marcos et al., 2014). In all the experiments, negative and positive controls were performed with distilled water and 2.5 mM methyl methane sulfonate (MMS), respectively. Three hundred eyes were scored per sex for every analysed solution or concentration. The number of hatched flies per bottle was used as a semiquantitative toxicity parameter, and the frequency of eyes with at least one mutant spot (mosaic eyes) was used to determine genotoxic activity.

With this assay, water samples and aqueous plant extract were analysed at different concentrations. Water samples were analysed directly as collected (UC, unconcentrated) and concentrated by lyophilization, moderately (MC; 3-4X, depending on the available sample) or highly (HC; 10X, for all the samples). After lyophilization, the samples were resuspended at the different concentrations with ultra-pure water. The aqueous plant extracts were prepared as described before (Sierra et al., 2025), although in some experiments the time of extract preparation was increased at 4, 8 and 24 h, to check its influence on the extract genotoxicity.

### 2.6. Genotoxicity Analysis: Comet Assay

The genotoxic activity of aqueous extracts from *Pteridium* plants was also analysed in human cells *in vitro*, with the comet assay (Collins 2004). HepG2 cells, from human hepatocellular carcinoma tumour cells, and HEK-293 cells, from human embryonic non-tumour kidneys, were cultured in Dulbecco’s Modified Eagle Medium (DMEM), with high glucose and L-glutamine (Biowest, France), supplemented with 10 % (v/v) FBS and 0,2 % PlasmocinTM Prophylatic (InvivoGen, EEUU), at 37 °C with 5% CO_2_ and 95 % relative humidity. For subculture, trypsin 1X was used at 80-90 % confluence. These cells were treated, in 6-well plates, with aqueous plant extract concentrations of 0.05, 0.1, 0.5 and 1 mg/mL, filtered through 0.2 μm syringe NalgeneTM filters for sterilization.

The comet assay was performed as described before (Espina et al. 2017; Rodríguez Pescador et al., 2022), with 10^5^cells per gel, in 0.5 % low melting point agarose gels, previously coated with 0.5 % normal melting point agarose. The gels were subjected to one hour of lysis (2.5 M NaCl, 0.25 M NaOH, 100 mM Na_2_EDTA, 10 mM Tris, pH 10, with 10% DMSO and 1% Triton X-100) at 4 °C, in darkness, 20 min of denaturation and 20 min of electrophoresis at 0.81 V/cm and 300 mA, both in 1 mM Na_2_EDTA and 300 mM NaOH pH13 buffer, at 4 °C. Then, the gels were neutralized with 0.4 M Tris buffer (pH 7.5), 3 times 5 min, fixed in absolute ethanol and dried at room temperature overnight. Slides were coded for blind analysis, and the gels were stained with 0.4 μg/mL ethidium bromide and Vectashield® fluorescence protector (VECTOR laboratories, Inc. Burlingame). An OlympusBX61 fluorescence microscope, with an Olympus DP70 digital camera, was used at 400X magnification to visualize the generated nucleoids and to take photos of at least 50 of them per gel. Photos were analyzed with the software program KOMET 5 (Kinetic Imaging Limited, now Andor-Oxford Instruments, Belfast, UK). The percentage of DNA in the comet tail (% Tail DNA) was the parameter used to measure DNA damage. Two gels were prepared per analyzed concentration in each experiment, and 3 independent experiments were performed for each sample in each cell line.

### 2.7. Allelopathic effects of bracken extract on the germination and growth of other species

To improve understanding of the potential allelopathic effects of *Pteridium aquilinum*, four common herbaceous species in Asturian grasslands were selected: *Trifolium repens, Bellis perennis*, *Agrostis capillaris*, and *Festuca pratensis*. The experiment consisted of germinating seeds of these four species in the presence of different concentrations of fern extracts.

For extract preparation, bracken samples collected in Somiedo (BRV) and Urbiés (MU11) were selected. The extracts were prepared in 250 mL glass flasks at a concentration of 0.5 g of lyophilized bracken samples/100 mL H₂O. The flasks were placed on an inversion shaker for 24 hours to extract the aqueous compounds. After 24 hours, the flasks were removed from the shaker and filtered. First, each extract was poured through a coarse sieve into beakers to remove large residues. Then, they were filtered again into new glass bottles using muslin with a pore size of 100–150 μm to eliminate finer particles.

Once the extracts were filtered, extract solutions at a final concentration of 5 mg/mL were labelled as “BRV 5” and “MU11 5”, and diluted solutions at final concentration of 1 mg/mL were marked as “BRV 1” and “MU11 1”. Distilled water was used as negative control. For germination experiments of each species and treatment, twenty-five seeds were placed on filter paper discs (Whatman No. 2, 70 mm diameter) moistened with 3 mL of the corresponding extract, or negative control, and placed in Petri dishes (90 mm diameter). A strip of absorbent paper soaked in water was also placed along the side of the dish to maintain humidity throughout the experiment. The dishes were carefully sealed with Parafilm and placed in a growth chamber at 25 ± 2 °C under a 16-hour photoperiod. Four replicates per treatment-species combination were used. The numbers of germinated seeds per extract and species were recorded every 2–3 days for one month. After one month, root length and number of leaves per seedling were recorded.

### 2.8. Statistical Analyses

The data presented in this work are either arithmetic means, with their standard deviations (SDs) or standard errors (SEs), or frequencies with their SDs, estimated as p·q/ √ N, where p is the estimated frequency of mosaic eyes, q is 1 − p, and N is the number of scored eyes (Marcos et al., 2014). Analysis of the frequencies of mosaic eyes were carried out with Chi-square tests. The results of hatched flies/bottle in the treatments were compared to those of the corresponding negative controls with Student’s *t* tests. Relationships among variables were evaluated by correlation and linear regression analyses. Metabolite contents were compared using Kruskal–Wallis tests followed by Dunn’s post hoc test. Due to non-normal distributions, differences between culture media were assessed using the Mann–Whitney U test, while the effects of culture medium, species, treatment, and their interactions were evaluated by two-way ANOVA with Tukey’s post hoc test. Statistical analyses were performed using IBM SPSS program (version 21.0.0.0) and R software (v. 4.4.1; R Core Team, 2024) using the Rcmdr package, with significance set at α = 0.05.

## 3. Results

### 3.1 In vitro spore culture of bracken

The fern life cycle comprises two heteromorphic phases: the gametophytic phase, which produces gametes, and the sporophyte phase, the dominant phase responsible for spore production. Observing these phases separately in nature is often challenging, making *in vitro* culture a valuable tool for their study (**Figure 4**). Sporulation of bracken is nearly absent in our region; however, in tropical climates, spore formation can occur year-round. In this study, spores from sporangia of plants growing in Honduras were successfully cultured *in vitro* (**Figures 4 a,b**). Following germination, gametophytes developed slowly over several weeks (**Figure 4c**), with gametangia, including both antheridia and archegonia, becoming visible under the microscope (**Figures 4 d,e**). When cultured on 1 MS medium, gametophytes formed aggregates that expanded rapidly through basal branching, yet sporophyte formation was absent (**Figure 4f**). In contrast, abundant sporophyte formation was observed on 1/2 MS medium (**Figure 4g**).

**Figure 4.**
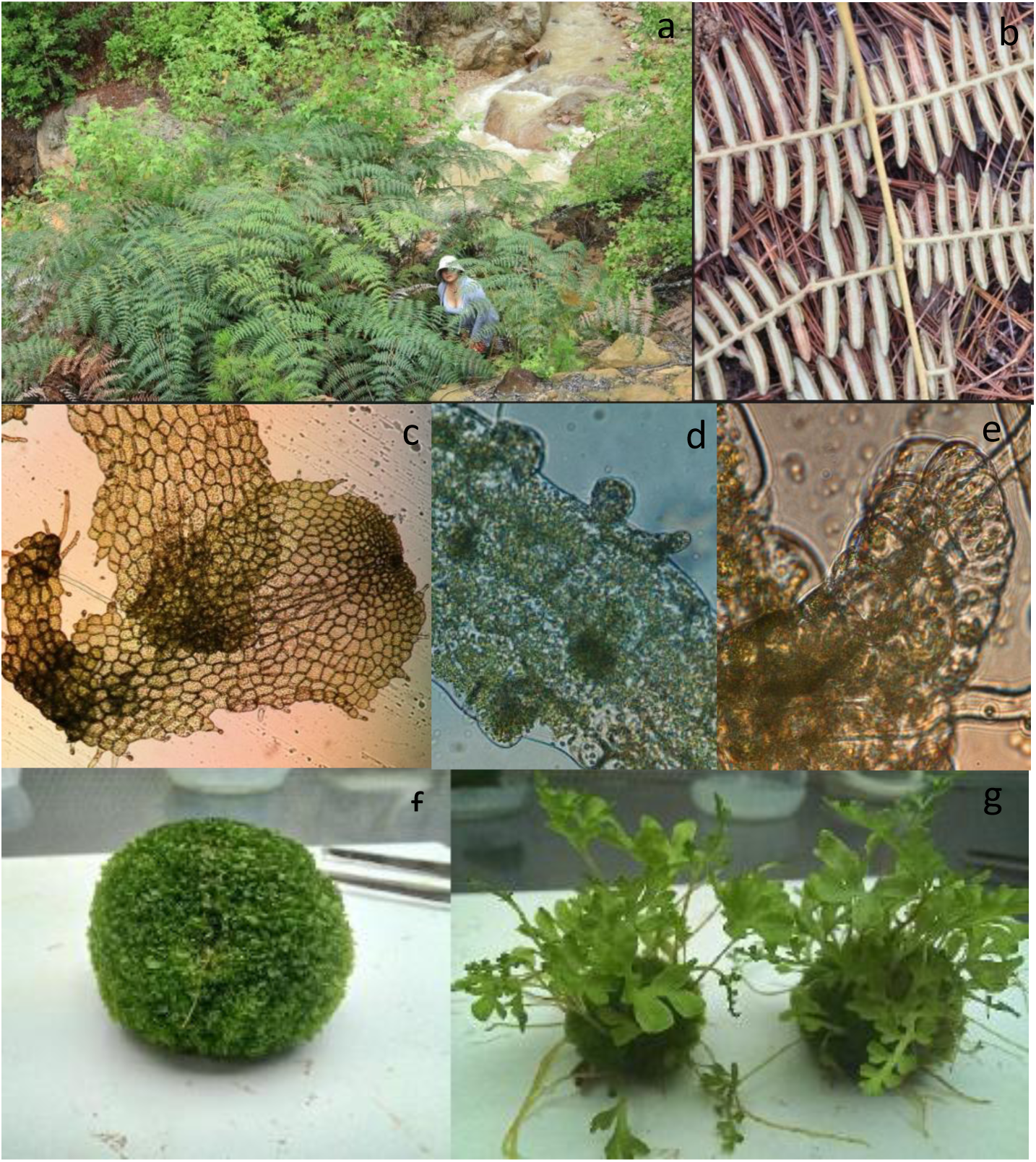
*In vitro* culture of *Pteridium aquilinum*. a) Tropical sporophytes in the field producing spores; b) Frond underside showing mature sporangia; c) Gametophyte growing *in vitro*; d, e) Detail of archegonia and antheridia; f, and g) Gametophyte clusters cultured on full, or half-strength MS culture media, respectively, forming abundant sporophytes in the diluted medium.

The effect of culture media, 1 and 1/2 MS, on sporophyte development was tested (**Figure 5**). In contrast to what occurs with the formation of sporophytes in the gametophyte, the number of fronds was favored in 1 MS medium (**Figure 5a**), while the frond length, and consequently the sporophyte length, showed no differences between the 1MS and 1/2 MS media (**Figure 5b**). The Mann-Whitney U test was used to analyse both parameters. Regarding the number of fronds per sporophyte, the test showed no significant difference between the groups (p= 0.1571). The W statistic was 2861.5, indicating that there is insufficient evidence to reject the null hypothesis. These results suggest that culture media do not significantly affect the number of fronds values. In relation to the length of sporophytes, the test revealed a statistically significant difference between the two groups (p< 0.0001). The median length of sporophytes was 2.5 for 1/2MS and 3.5 for 1MS, indicating that 1MS supports significantly longer sporophytes. These results suggest that the medium type has a significant effect on sporophyte growth.

**Figure 5.**
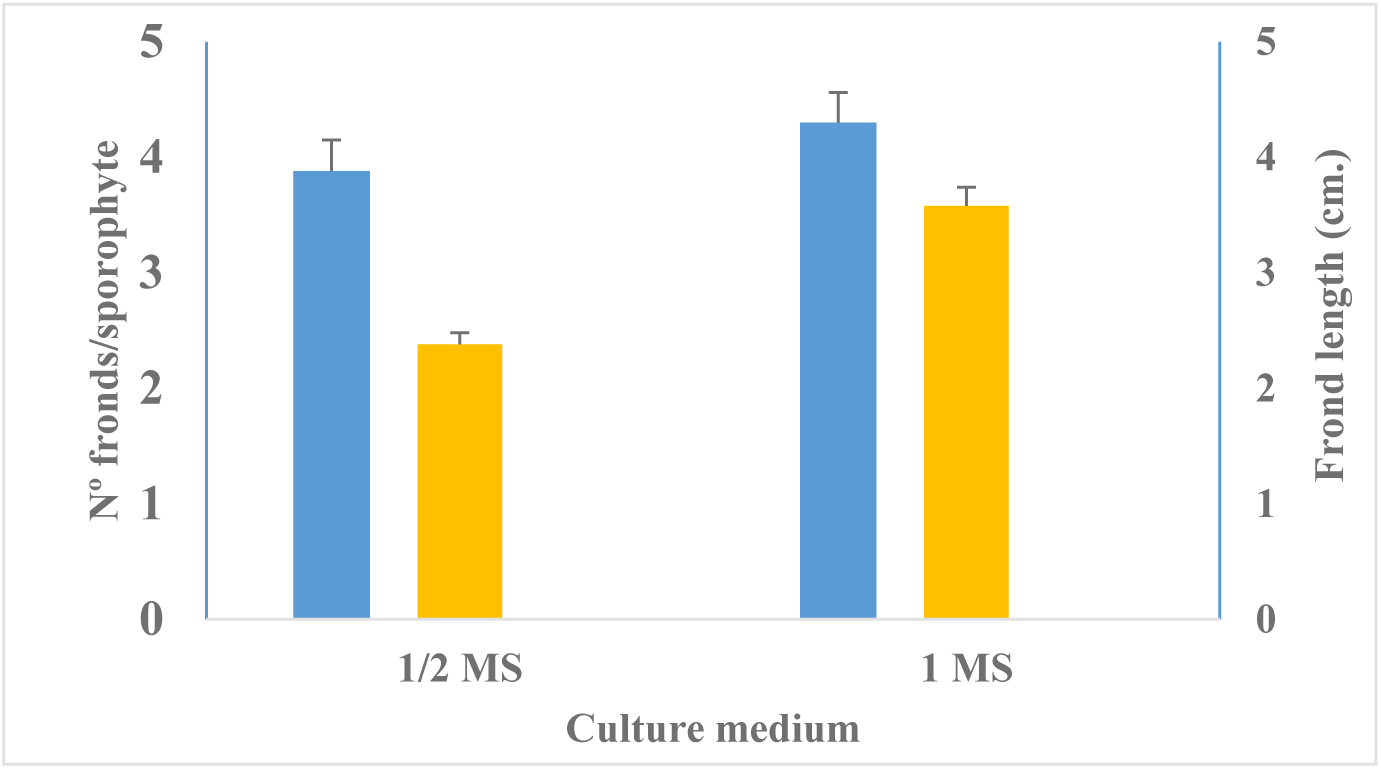
Growth of sporophytes of *Pteridium aquilinum*, cultured in full and half-strength MS medium, estimated as: a) number of fronds per gametophyte cluster, and b) length of fronds. Data are presented as arithmetic means with their standard errors.

### 3.2 Purification tracking by HPLC-PAD

Following extraction and filtration, HPLC–PAD analysis of the extract revealed chromatographic signals whose retention times corresponded to those of pterosins Ptr B and Ptr A, confirmed by direct comparison with commercial standards, and a third one which was tentatively assigned to Ptr G (**Figure 6**). After subjecting the extract to hydrolysis, the marked increase in the intensity of three chromatographic signals reinforced this assignment by its UV–visible absorption spectrum (217, 263, and 306 nm), and elution order. Thus, it is consistent with previous reports indicating that the studied plant species contains high levels of PTE, whose hydrolysis leads to the formation of Ptr G (Kisielius et al., 2020).

**Figure 6.**
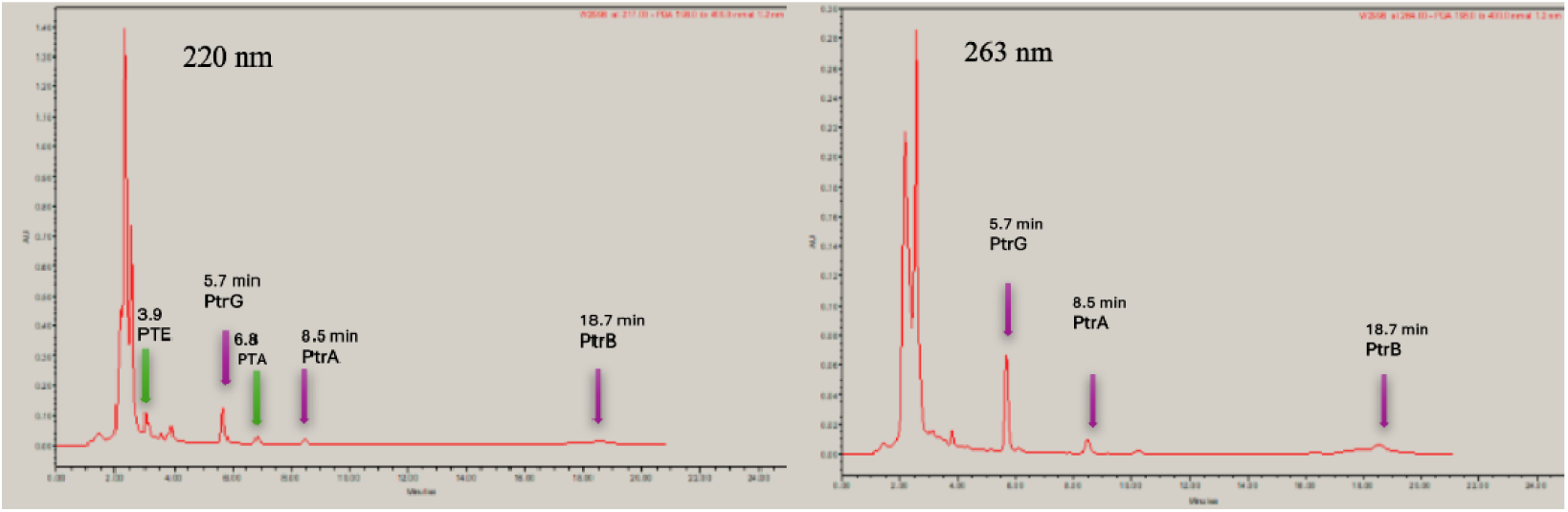
Chromatogram of the aqueous extract of *Pteridium esculentum* at 220 nm and 263 nm.

Subsequent passage of the extract through a polyamide resin resulted in the disappearance of the signals corresponding to free pterosins (**Figure S2**). To facilitate the identification of glycosylated derivatives, a fraction of the extract was hydrolyzed and the chromatographic profiles before and after hydrolysis were compared, revealing the disappearance of certain signals and the concomitant increase in others.

Based on this comparative analysis, the extract was further purified using an SPE-HLB cartridge. HPLC–PAD analysis of the resulting fraction revealed three well-resolved signals with retention times of 3.87, 5.43, and 6.91 minutes (**Figure S3**), whose UV–visible absorption spectra were consistent with those reported for illudane glycosides in the literature (Kisielius et al., 2020).

### 3.3 Chromatographic fractionation and identification by mass and ¹H NMR analysis. Determination of compounds purity

The extract processed by the polyamide-SPE HLB was analyzed for its respective fractionation in the HPLC-DAD Agilent equipment, obtaining the results presented on **Figure 7**.

**Figure 7.**
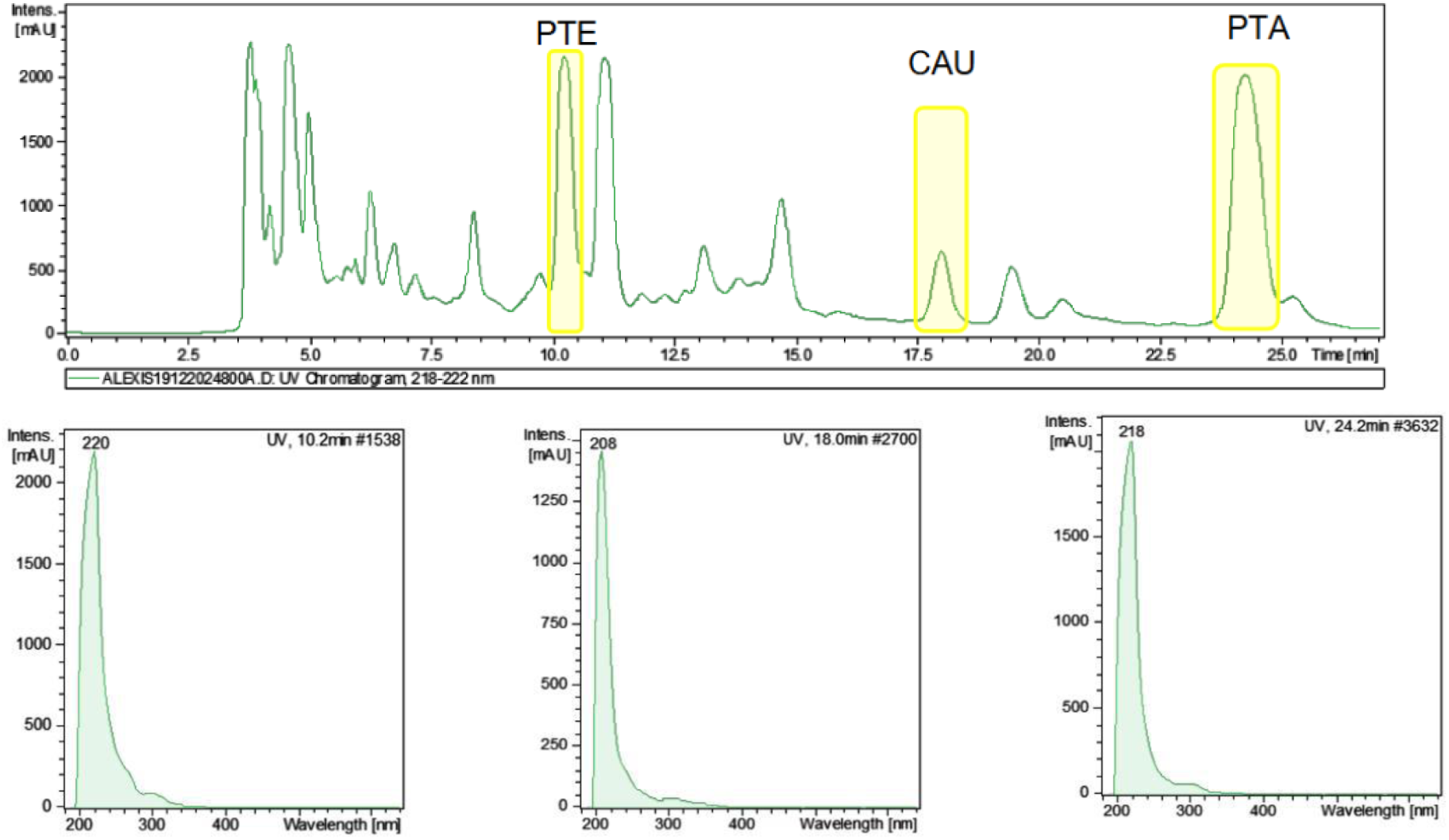
HPLC-DAD Agilent chromatogram of extract processed by polyamide

Once signals with absorption spectra matching those reported in the literature for illudane glycosides were identified, the corresponding fractions were isolated and grouped as R01, R02, and R03. These fractions were subsequently frozen and lyophilized to obtain solid residues. In the case of R03, rapid rehydration of the solid was observed, indicating hygroscopic behaviour. Finally, three glycosylated illudanes were identified in fractions R01, R02, and R03 isolated from Pteridium using HPLC–PDA, UPLC–HRMS, and ^1^H NMR analyses (**Figure S4**). Fraction R01 showed UV absorption at 203 nm and characteristic UPLC–HRMS ions consistent with ptesculentoside (C₂₀H₃₀O₉Na), which was confirmed by ^1^H NMR and showed 83.10% purity (**Figures S5–S10**, **Table S1**). Fraction R02 displayed UV absorption at 204 nm and diagnostic UPLC–HRMS ions corresponding to caudatoside (C₂₁H₃₂O₉Na), with identity confirmed by ^1^H NMR and a purity of 66.32% (**Figures S11–S14, Table S2**). Fraction R03 exhibited UPLC–HRMS ions consistent with ptaquiloside (C₂₀H₃₀O₈Na), which was confirmed by ^1^H NMR and showed 93.33% purity (**Figures S15–S18, Table S3**).

Overall, the results demonstrate the simultaneous occurrence of PTE, CAU and PTA in the evaluated fractions and highlight the robustness of the combined chromatographic–mass spectrometric–NMR approach for illudane structural characterization in Pteridium. However, the purity of CAU was insufficient to employ it as a reference standard for quantitative analysis. Consequently, quantitative determination was performed exclusively using PTA, PTE, PtrB, and PtrA as analytical standards.

### 3.4. Quantification of PTA, PtrB, Ptr A and PTE

The results obtained from the analysis of the four compounds: PTA and corresponding PtrB, and also PtrA, a metabolite derived from CAU, and the illudane glycoside PTE are shown in **Figure 8**. All four compounds have been detected in all the analyzed plant samples. The results of the Kruskal-Wallis test indicated significant differences in the concentrations of the metabolites PTE, PTA, PtrA, and PtrB between the different plant tissues (p< 0.05 for all metabolites), suggesting that the distribution of metabolites varies significantly by plant type. To perform post-hoc comparisons, the Dunn’s test was applied, revealing significant differences in some comparisons. Specifically, for PTE and PTA, BRV showed a significant difference with gametophyte (p= 0.0016). In the case of PtrA, significant differences were observed between PEBEL and BRV (p= 0.0031), and between BRV and gametophyte (p= 0.0256). Similarly, for PtrB, comparisons between PEBEL and BRV were also significant (p= 0.0031). Adjusted p-values were used to account for multiple comparisons.

**Figure 8.**
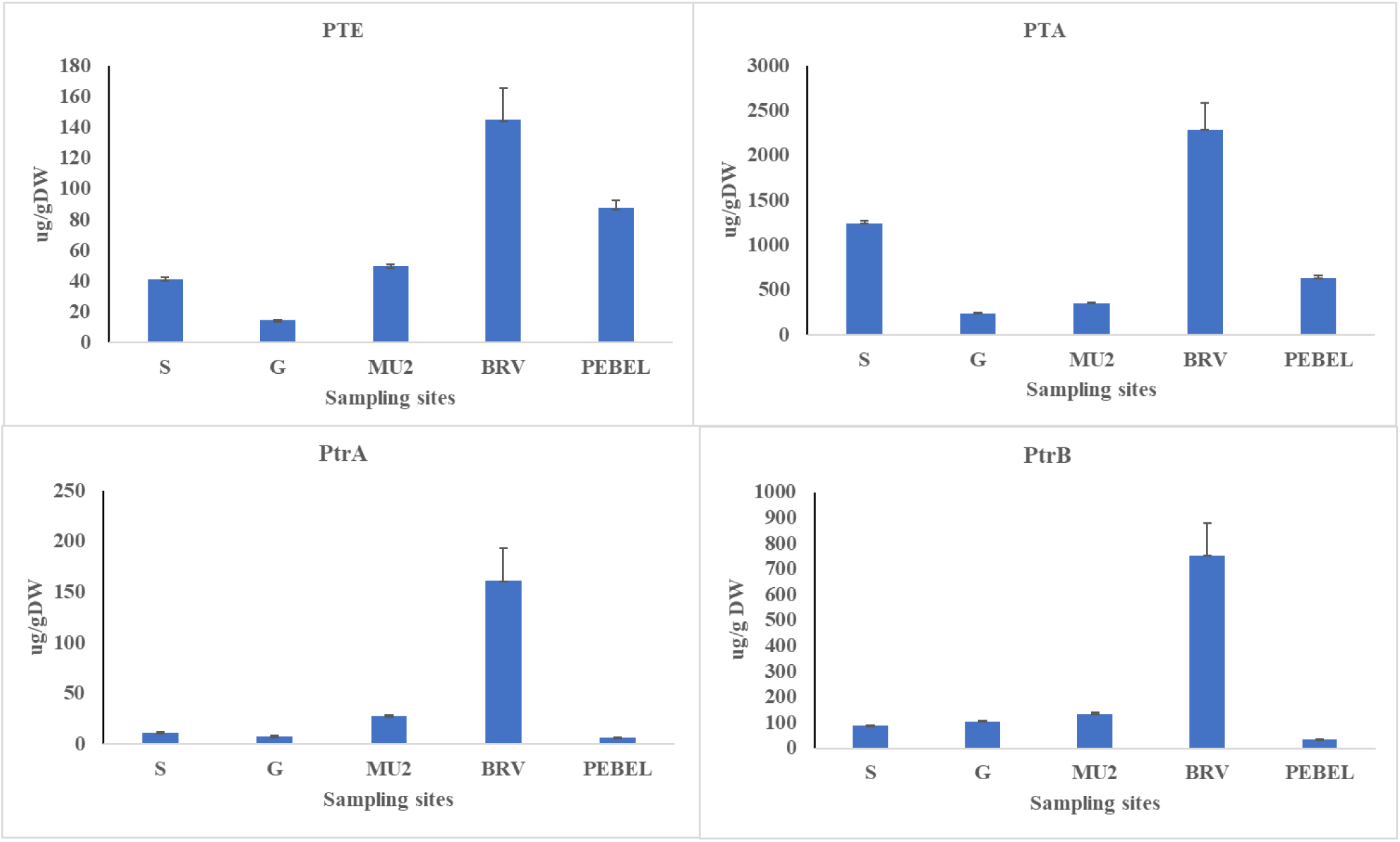
Levels of the illudane glycosides PTA, PTE and of the pteriosins PtrA and PtrB in sporophyte (S) and gametophyte (G) of bracken derived from spores collected in Honduras and then cultured *in vitro*, and in croziers gathere in spring in three spots of Asturian region: Montes de Urbiés (MU2), and the Natural Parks of Somiedo (BRV), and Picos of Europe (PEBEL).

Strikingly high levels of PTA were detected in the BRV site of the Somiedo Natural Park, where there were records of cow deaths, and which reached orders of milligram per gram of plant dry weight. Likewise, the highest levels of the rest of the metabolites analysed were also found in this same area. Although no CAU pattern was available, the levels of its associated metabolite PtrA indicated CAU presence in all the populations studied, and especially in the BRV sample. Moreover, levels of PTE, which is less frequent in European latitudes, were also detected. On the other hand, the levels of PTA and PTE detected in the sporophyte from Honduran spores cultured *in vitro*, and of PTE present in the populations of the Picos de Europa PEBEL spot, were strikingly high.

Regarding the levels of metabolites analysed in the water samples, from ten areas of the Asturian geography (**Table 1**), the results obtained, presented in **Figure 9**, revealed the presence of PTA in 4 of them: MU2, MU11, PON and RED; being the highest detected in PON. On the other hand, the two pterosins analysed were detected in all the samples, suggesting the presence of CAU, and of PTA, in all studied areas, even though this last chemical was not directly detected. Finally, the illudane glycoside PTE was only absent in PRI and DEG. Regarding metabolite concentrations, PTA reached up to 1,6 µg/L at RED, PtrB up to 6,3 µg/L, PtrA up to 2,8 µg/L, and PTE up to 2,6 µg/L.

**Figure 9.**
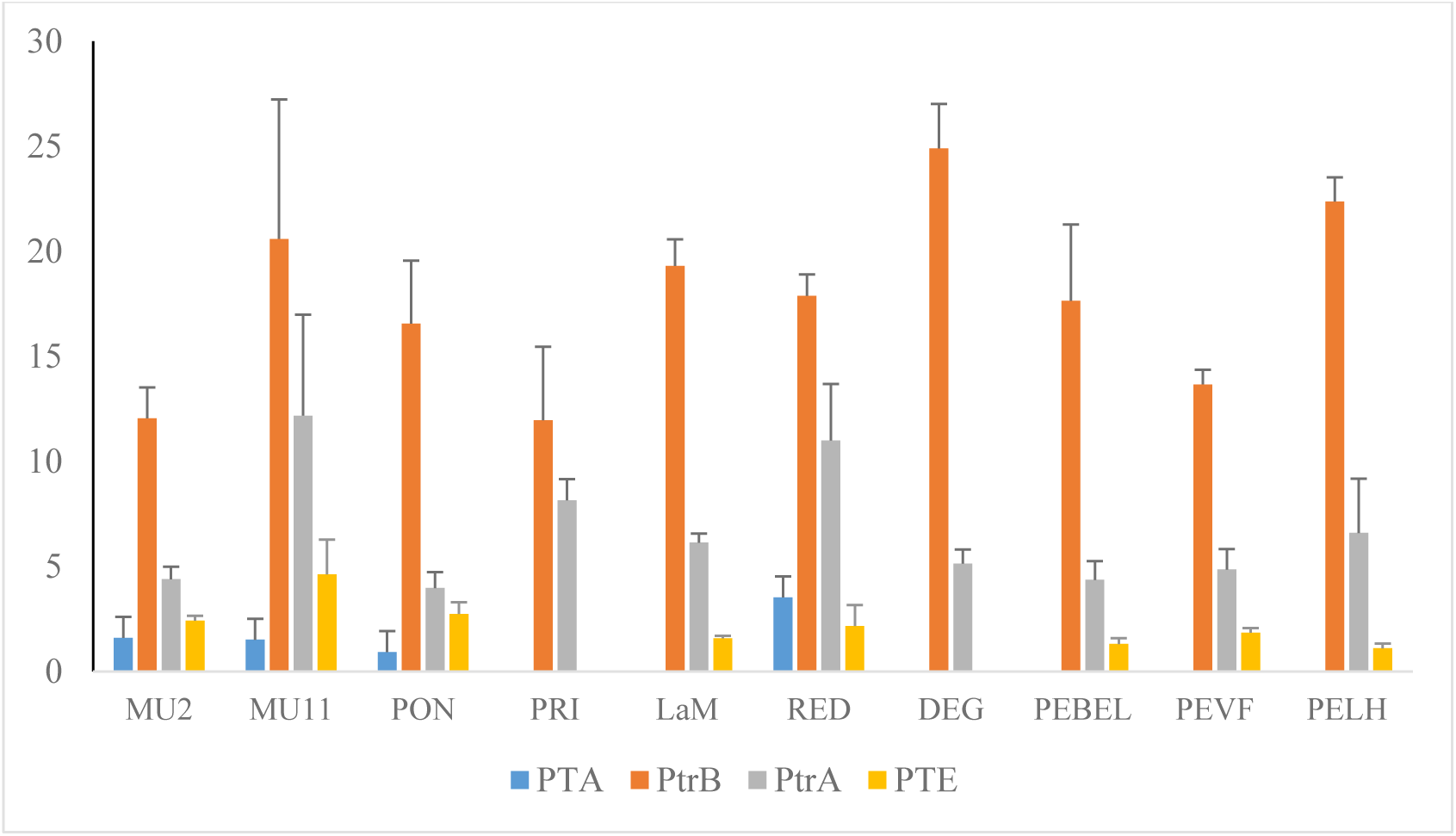
Levels of the illudane glycosides PTA, PTE and the pteriosins PtrA and PtrB in samples of water sources collected in several areas infested with bracken. Data are presented as arithmetic means with their standard errors.

The Kruskal-Wallis test did not indicate any significant differences between the locations for PTA, PTE and PtrB. For PtrA, although the Kruskal-Wallis test was significant, Dunn’s test did not reveal any pairwise differences between the locations after adjusting for multiple comparisons. Overall, these findings indicated that there were no differences in metabolite concentrations across the groups in this dataset.

### 3.5. Genotoxicity analyses of water samples

Among all the water samples collected in this work, genotoxic activity was determined *in vivo* for 4 of them: MU2, MU11, PEBEL (BEL) and BRV, with the SMART assay. Results of toxicity and genotoxicity are presented in **Figure 10**. First, none of the samples induced toxicity. With respect to genotoxic activities, the highest concentration of all the samples (HC) induced frequencies of mosaic eyes higher than the respective spontaneous ones, both in females and males. The intermediate concentration (MC) induced frequencies of mosaic eyes higher than the spontaneous ones with all the samples but BRV in males, and with only MU2 and MU11 in females. The unconcentrated samples did not induce frequencies of mosaic eyes over the negative controls. Moreover, the comparisons between females and males revealed that, contrary to what was described for aqueous plant extract, the water samples did not induce significative recombination, as observed in **Figure S18.**

**Figure 10.**
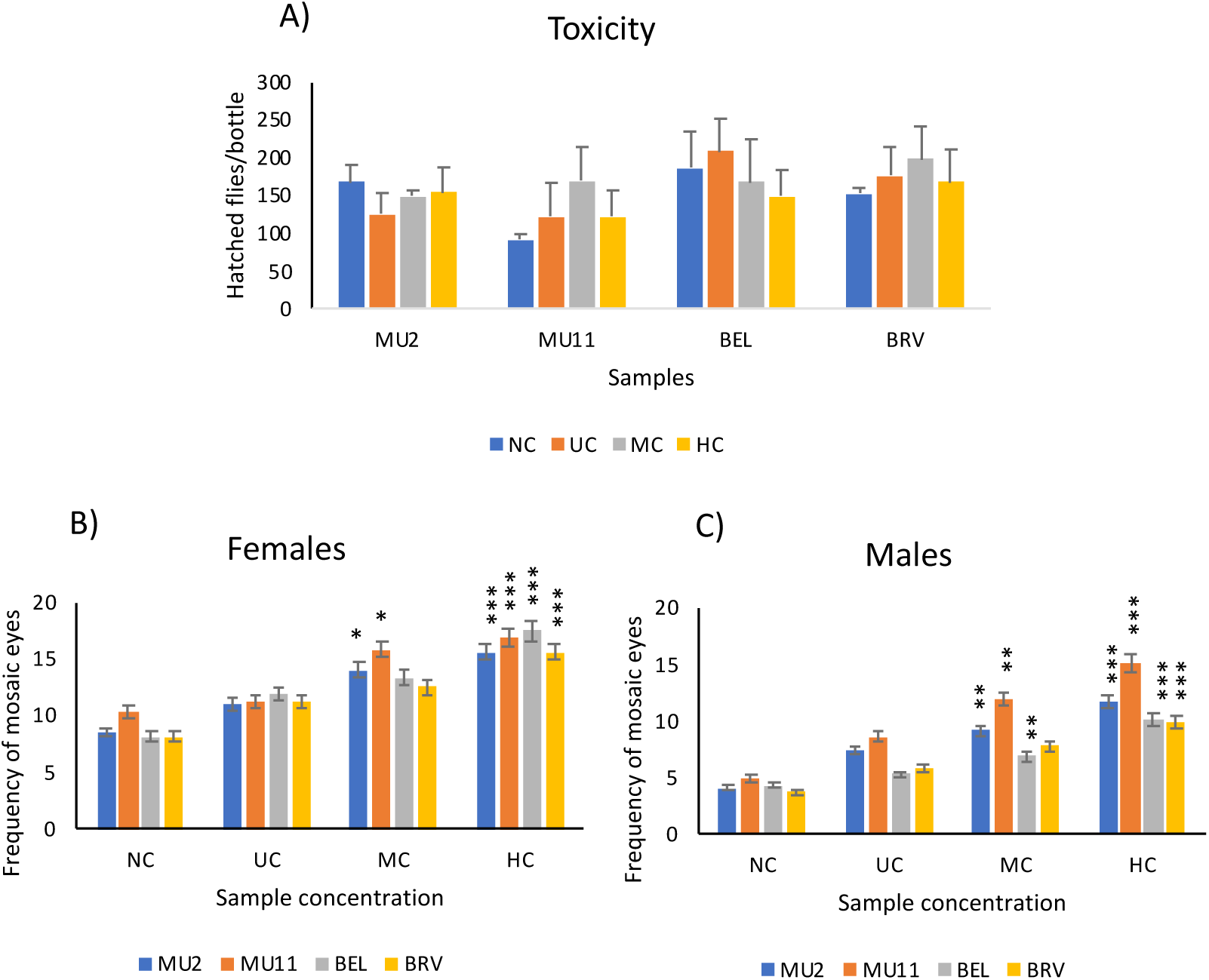
Toxicity and genotoxicity analysis of water samples. A) Hatched flies per bottle as a measured of toxicity, numbers are arithmetic means of at least 4 bottles, with the corresponding standard errors. Frequencies of mosaic eyes in negative controls (NC) and unconcentrated (1x or UC), moderately concentrated (3-4X or MC) and highly concentrated (10x or HC) treatments, B) in females and C) in males. Values are frequencies of at least two independent experiments and their corresponding standard deviations. Comparisons between treatments and the corresponding negative controls: * p< 0.05; ** p< 0.01; *** p< 0.001.

The frequencies of mosaic eyes induced by these samples were compared to the levels of metabolites described in **Figure 9**. Results revealed that there was no significant relationship between these two parameters. However, when these frequencies of induced mosaic eyes were compared to those induced by aqueous extracts obtained from plants collected at the same places, significant correlations were found, both in females (R= 0.885, p< 0.001, 10 d.f.) and in males (R= 0.60, p< 0.05, 10 d.f.), when all the concentrations from all the samples were included in the analysis.

### 3.6. Genotoxicity analyses of aqueous plant extracts

In this work, to finish the genotoxic analysis of plant samples collected in Asturias, a new sample BRV, not included in our previous work (Sierra et al., 2025), was chosen because the collecting site was related to several cattle deaths. Results of this analysis are presented in **Figure 11**. All the extract concentrations increased the frequencies of mosaic eyes over that of the negative control in females, and all but the lowest concentration did it in males.

**Figure 11.**
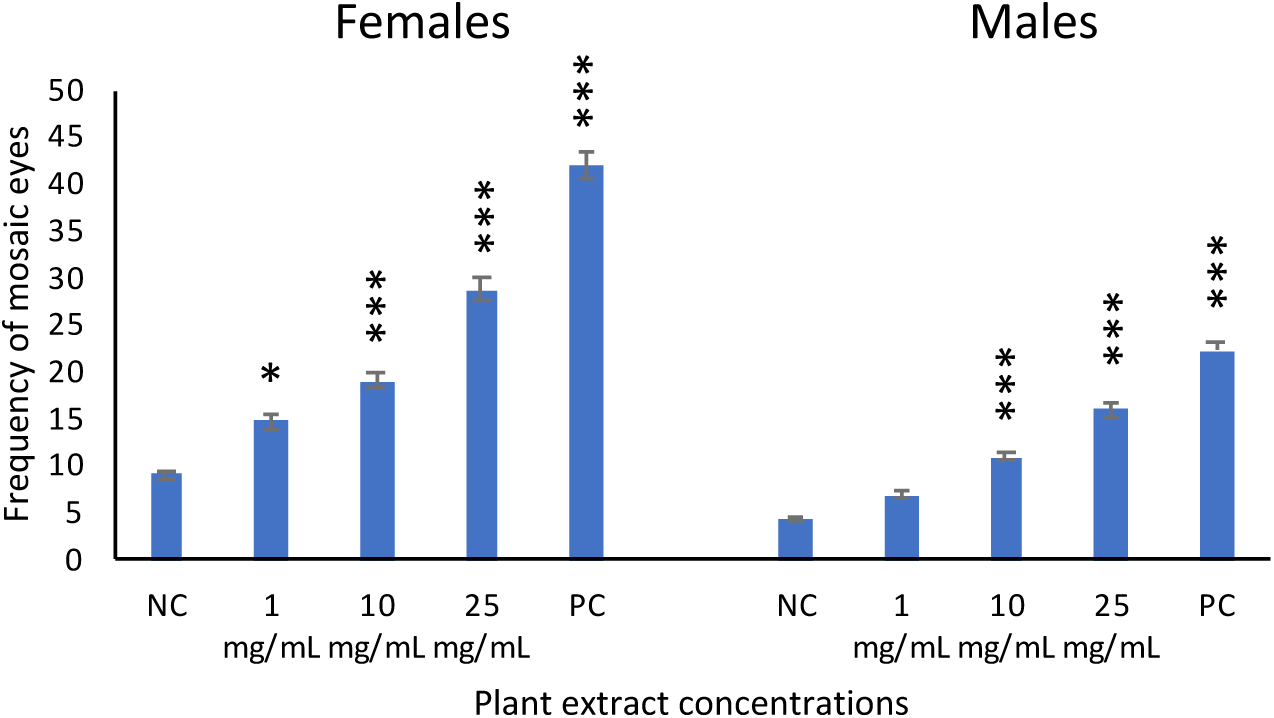
Genotoxic analysis of BRV plant extracts. Frequency of mosaic eyes in negative and positive controls (NC and PC, respectively), as well as in the treatments with the different plant extract concentrations. Comparisons between treatments and the corresponding negative controls: * p< 0.05; *** p< 0.001.

When these results were joined with those already published, the relationship with latitude, described before (Sierra et al., 2025), disappeared. At this point, since there were metabolite data for all the samples analyzed at the genotoxicity level (17 samples), it was possible to confirm the maintenance of the statistically significant correlations between the induced frequencies of mosaic eyes and the levels of pterosin B in females (with 15 d.f., R= 0.71, p< 0.01, R= 0.79, p< 0.001, R= 0.62, p<0.01, respectively for 1, 10 and 25 mg/mL concentrations) and in males (with 15 d.f., R= 0.51, p< 0.05, R= 0.635, p< 0.01, R= 0.60, p< 0.05, respectively for 1, 10 and 25 mg/mL concentrations). The statistically significant correlation with the levels of pterosin A was also maintained in females (with 15 d.f., R= 0.65, p< 0.01, R= 0.65, p< 0.01, R= 0.48, p< 0.05, respectively for 1, 10 and 25 mg/mL concentrations), and even increased in males because a significative relationship was also found for 1 mg/mL concentration (with 15 d.f., R= 0.56, p< 0.05, R= 0.64, p< 0.01, R= 0.54, p< 0.05, respectively for 1, 10 and 25 mg/mL concentrations).

In addition to this analysis, the SMART assay was also used to determine the effects of frond senescence on the genotoxic activities of aqueous extracts. For that, aqueous extracts from young fronds, collected in April, were compared to those from senescent fronds, collected in November, at MU2 and MU11 sites. Results, presented in **Figure 12**, showed that the frequencies of mosaic eyes induced by extracts from old fronds in females were significantly lower than those induced by extracts from young ones, although they were still higher than the corresponding spontaneous frequencies, at least with the highest analyzed concentration (25 mg/mL). On the contrary, in males, the frequencies of mosaic eyes induced by extracts from old fronds were not only lower than those induced by extracts from young ones, but they were not different from the corresponding spontaneous frequencies in any case.

**Figure 12.**
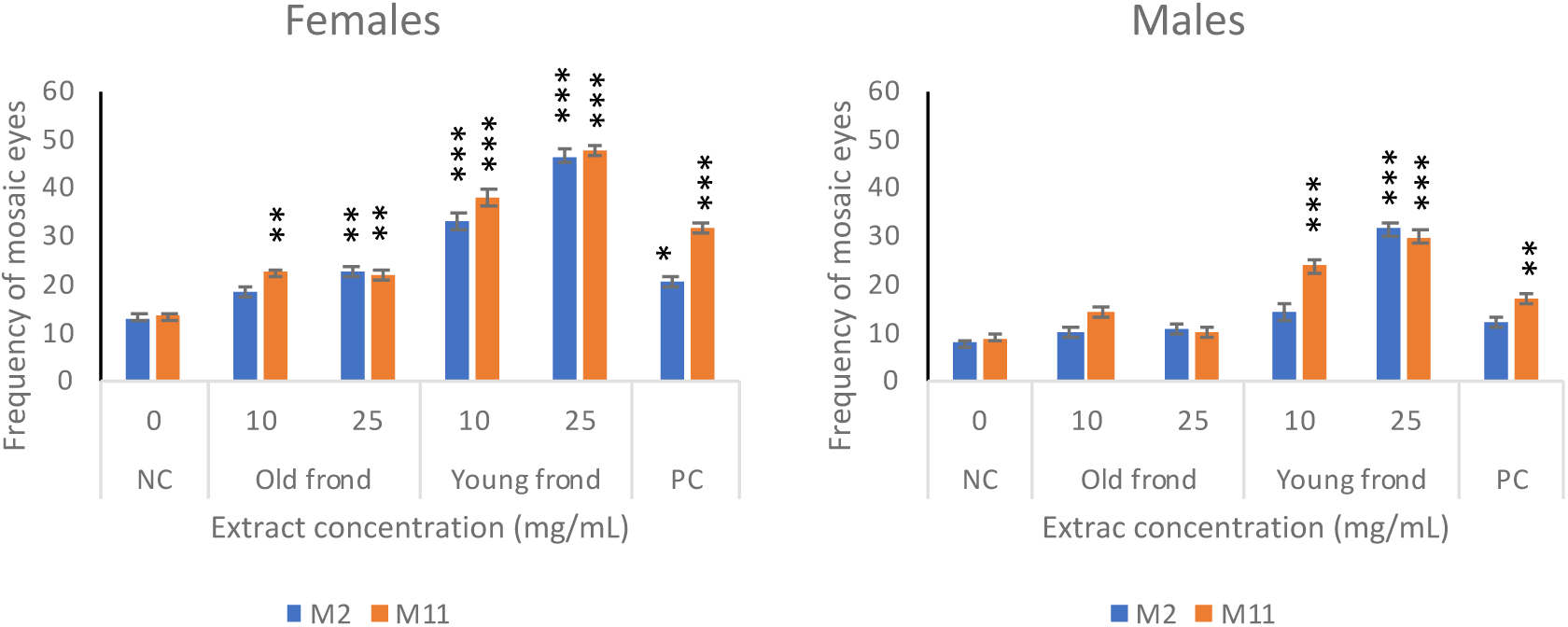
Effect of senescence on genotoxic activity. Comparison between the mosaic eye frequencies induced by young and senescent fronds from the MU2 and MU11 samples, in both females and males. Extract concentrations of 10 and 25 mg/mL have been analyzed. 2.5 mM MMS has been used as a positive control. * p< 0.05; ** p< 0.01; p< 0.001 compared to the respective negative controls (NC).

Moreover, this assay was also used to determine whether the time of the aqueous extraction influenced the genotoxic activity measured as frequency of mosaic eyes. In addition to 2.5 h, longer times, or 4, 8 and 24 h, were also studied with MU2 and MU11 samples. The obtained results, shown on **Figure 13**, demonstrated that, independently of the sample, mosaic eye frequencies increased with the extraction time up to 8 h, especially in females. Extracts obtained after 24 h of continuous agitation, by vertical rotation, induced mosaic eye frequencies, significantly higher than the respective spontaneous ones, but lower than those generated by 8 h extracts. According to these results, it was evident that the extraction time used in all the work carried out so far, of 2.5 h, was a good choice, and the most appropriate one for practical reasons of organization in the laboratory.

**Figure 13.**
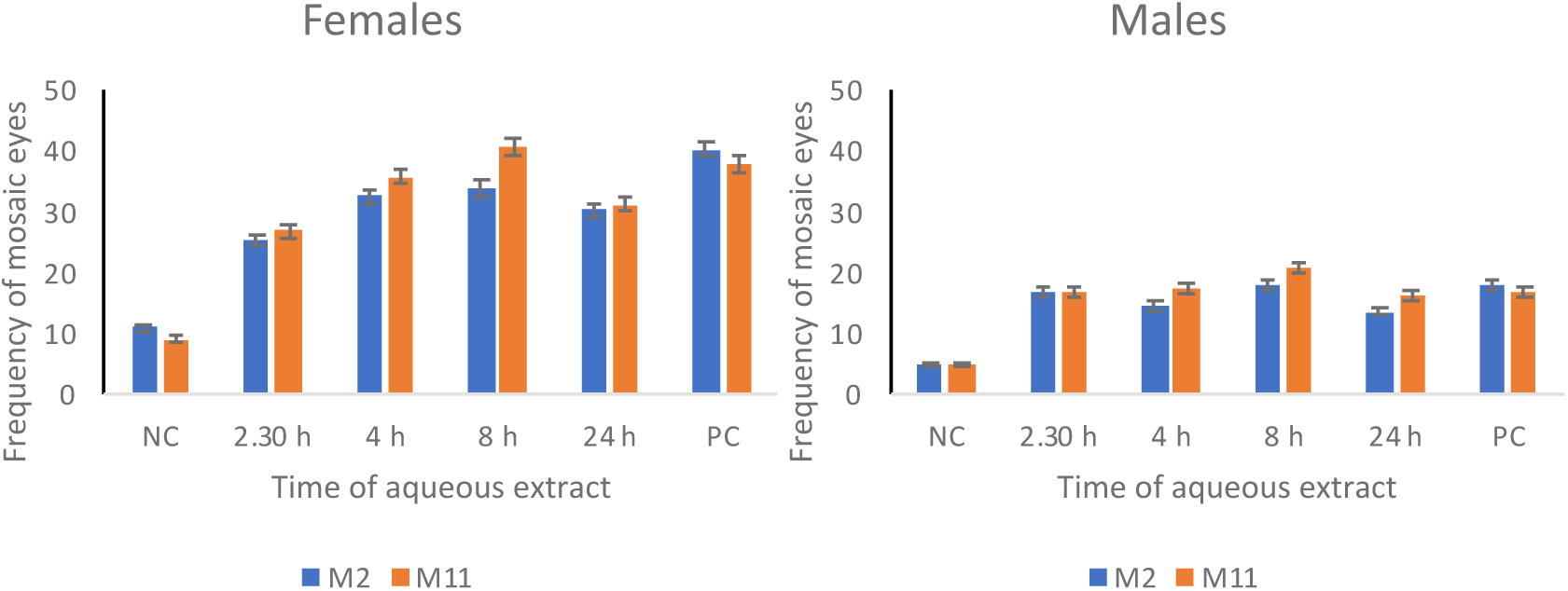
Extraction time influences the genotoxic activity of the samples. Frequencies of mosaic eyes induced by aqueous plant extracts, from MU2 and MU11 samples, generated with 10 mg/mL concentration and different extraction times. 2.5 mM MMS has been used as a positive control. All the times analyzed, for the two samples, induced mosaic eye frequencies statistically higher than their respective negative controls, with p< 0.001.

The genotoxic activities of aqueous extracts from Pteridium plants were also studied in human cells, *in vitro*, with the comet assay (Collins 2004). HepG2 and HEK-293 human cells, tumor and non-tumor ones, respectively, were treated with the MU11 and TIN samples (described in Sierra et al., 2025), and the results are presented in Figure 14.

**Figure 14.**
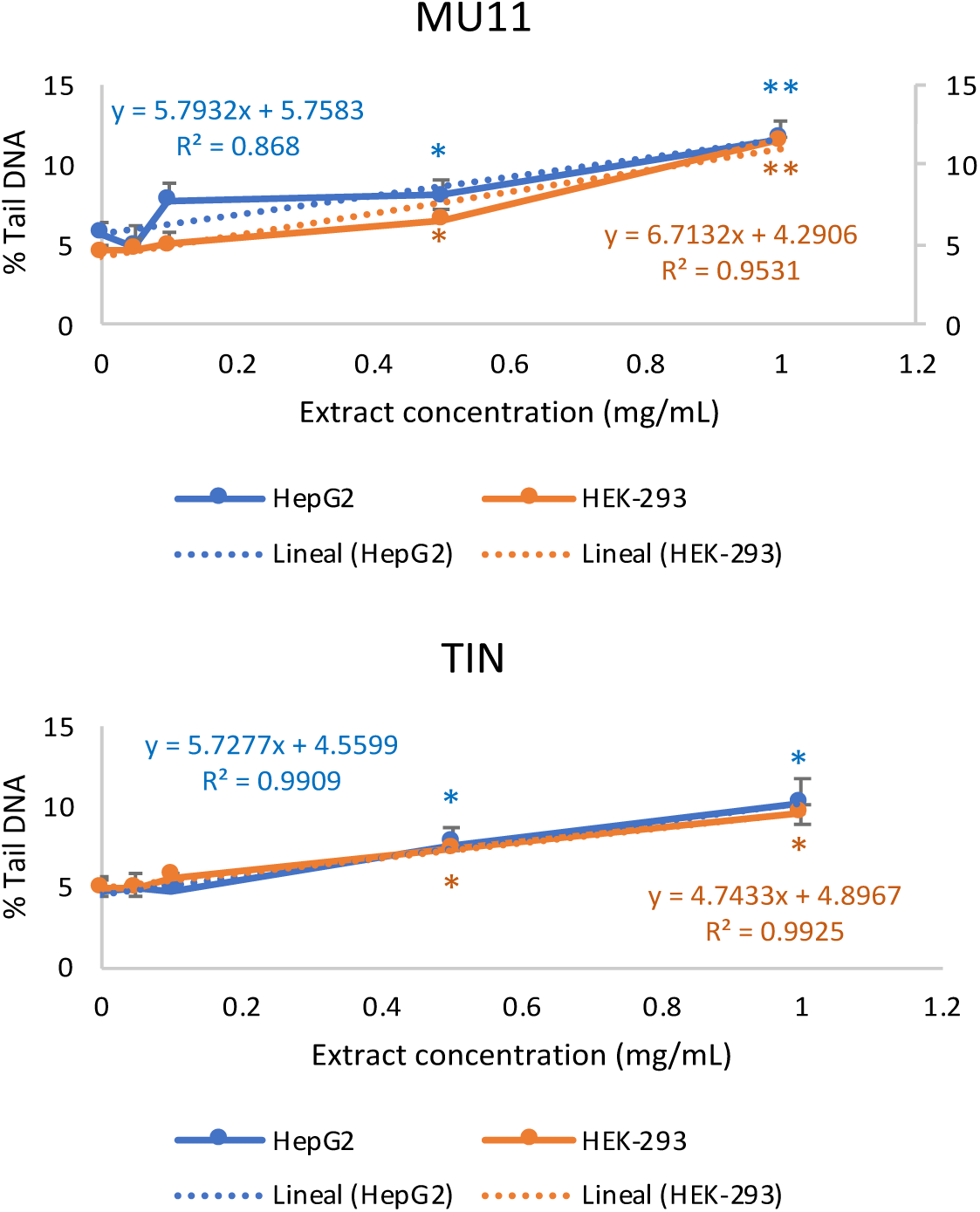
Analysis of DNA damage induced by bracken aqueous extracts, using the comet assay. Percentages of tail DNA induced with aqueous extracts from MU11 and TIN samples in tumor HepG2 and non-tumor HEK-293 cells. The regression lines are presented with their corresponding equations. * p< 0.05, ** p< 0.01 compared to the respective negative controls.

Results demonstrated that extracts from both samples induced percentages of tail DNA higher than the respective spontaneous ones with the highest analyzed concentrations, in both cell types. In addition, the extracts induced DNA damage that increased with concentration, as indicated by the dose-response regression analysis, with slopes significantly higher than zero in all cases (b= 5.79 ± 1.30, p= 0.019 for MU11-HepG2; b= 6.71 ± 0.86, p= 0.003 for MU11-HEK-293; b= 5.73 ± 0.32, p=0.0002 for TIN-HepG2; b= 4.74 ± 0.24, p= 0.00017 for TIN-HEK-293). Moreover, these data showed that the two samples induced equivalent damage in HEK-293 cells, but MU11 seems to induce slightly more DNA damage than TIN in HepG2 cells.

### 3.4 Allelopathic effects on germination and growth of herbaceous species

Seed germination of four plant species was evaluated under five different treatments (Control, BRV1, BRV5, M11 1, and M11 5) with four replicates per treatment, and results are shown in Figure 15. A two-way ANOVA (Species × Treatment) revealed statistically significant differences in germination among species (F₃,₆₀ = 149.95, p< 0.001) and in species × treatment interaction (F₁₂,₆₀ = 2.67, p= 0.006), but not in germination (F₄,₆₀ = 2.39, p= 0.061). Mean germination values by species showed numerical differences among treatments: *Agrostis capillaris* exhibited higher germination under M11 1 (81.9%) and M11 5 (73.0%) compared to the negative control (71.0%), while lower under BRV1 and BRV5 (59.0%). In *Bellis perennis*, BRV1 (84.0%) and M11 1 (89.4%) showed higher germination values relative to the control (78.1%). *Festuca pratensis* generally exhibited low germination, with slight increases under M11 1 (36.0%) and M11 5 (36.0%) compared to the control (21.0%). *Trifolium repens* showed the highest germination under BRV5 (90.0%) and BRV1 (86.0%), while the control reached 75.0%. Although no post-hoc comparisons were statistically significant, these results indicated species-specific responses to treatments, highlighting the importance of the species × treatment interaction when evaluating bracken treatment effects on seed germination. Overall, there was no clear evidence of allelopathic inhibition, but some treatments numerically increased germination in certain species.

**Figure 15.**
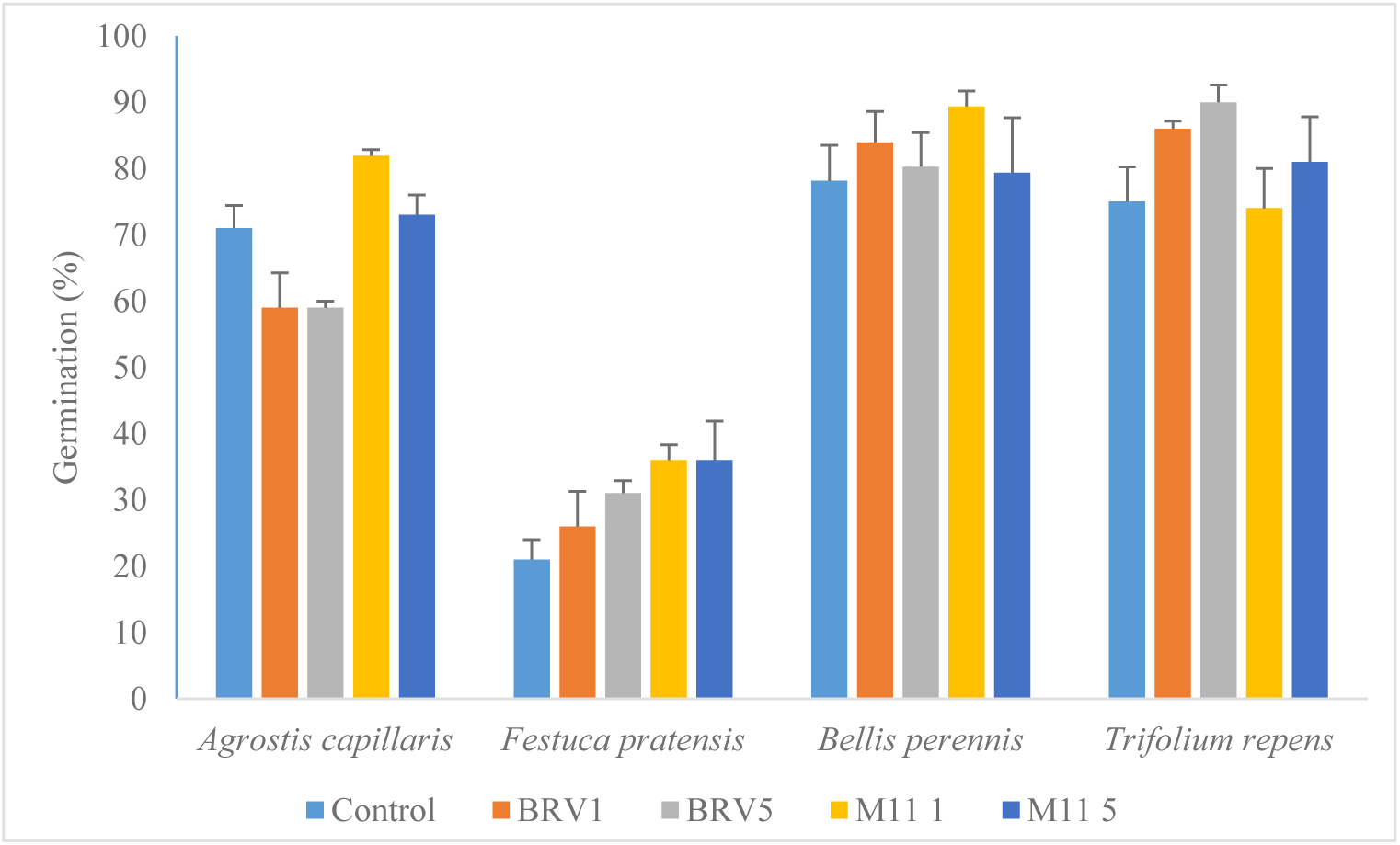
Effect of aqueous extracts on germination. Germination of four herbaceous seeds under exposition to aqueous extracts from Pteridium fronds. Data are presented as arithmetic means with their standard errors.

The effect of species and treatment on number of leaves and root length was evaluated using a two-way ANOVA (**Figures 16 and 17**). Both traits were significantly affected by species (root length: F₃,₁₁₇₂ = 192.17, p< 0.001; leaf number: F₃,₁₁₁₂ = 103.72, p< 0.001) and by treatment (root length: F₄,₁₁₇₂ = 7.25, p< 0.001; leaf number: F₄,₁₁₁₂ = 12.00, p< 0.001). Significant species × treatment interactions were also detected for both variables (root length: F₁₂,₁₁₇₂ = 3.87, p< 0.001; leaf number: F₁₂,₁₁₁₂ = 3.39, p< 0.001), indicating that the effect of treatments on early growth was species-specific. Mean comparisons showed clear differences among treatments within each species. *Agrostis capillaris* developed its longest roots under BRV 1 and M11 5 and produced the highest number of leaves under BRV 5. In *Bellis perennis*, root length increased under BRV 5 and M11 5, and leaf number was also higher under these treatments. *Festuca pratensis* showed its greatest root length and leaf number under BRV 1. *Trifolium repens* exhibited moderate increases in both traits, with roots being longest under M11 5 and leaf number highest under BRV 5. Overall, these results indicate that both root development and leaf production were positively influenced by the treatments, with no evidence of inhibitory effects on early plant growth. The significant species × treatment interactions highlight the importance of species-specific responses when evaluating the effectiveness of bracken treatments on plant performance.

**Figure 16.**
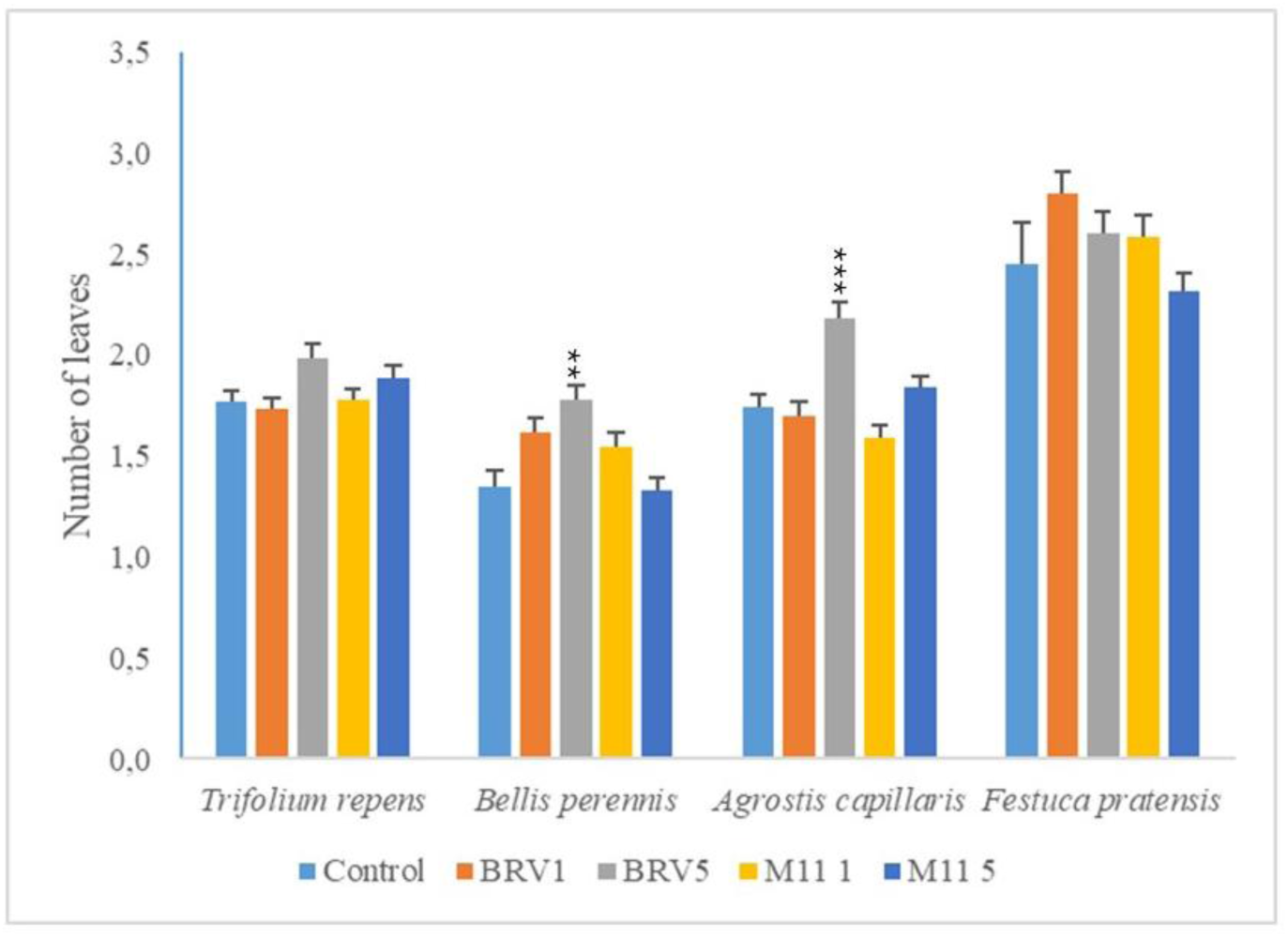
Effect of aqueous extracts on plant leaves. Number of leaves of the different species in the different culture media. Data are presented as arithmetic means with their standard errors.

**Figure 17.**
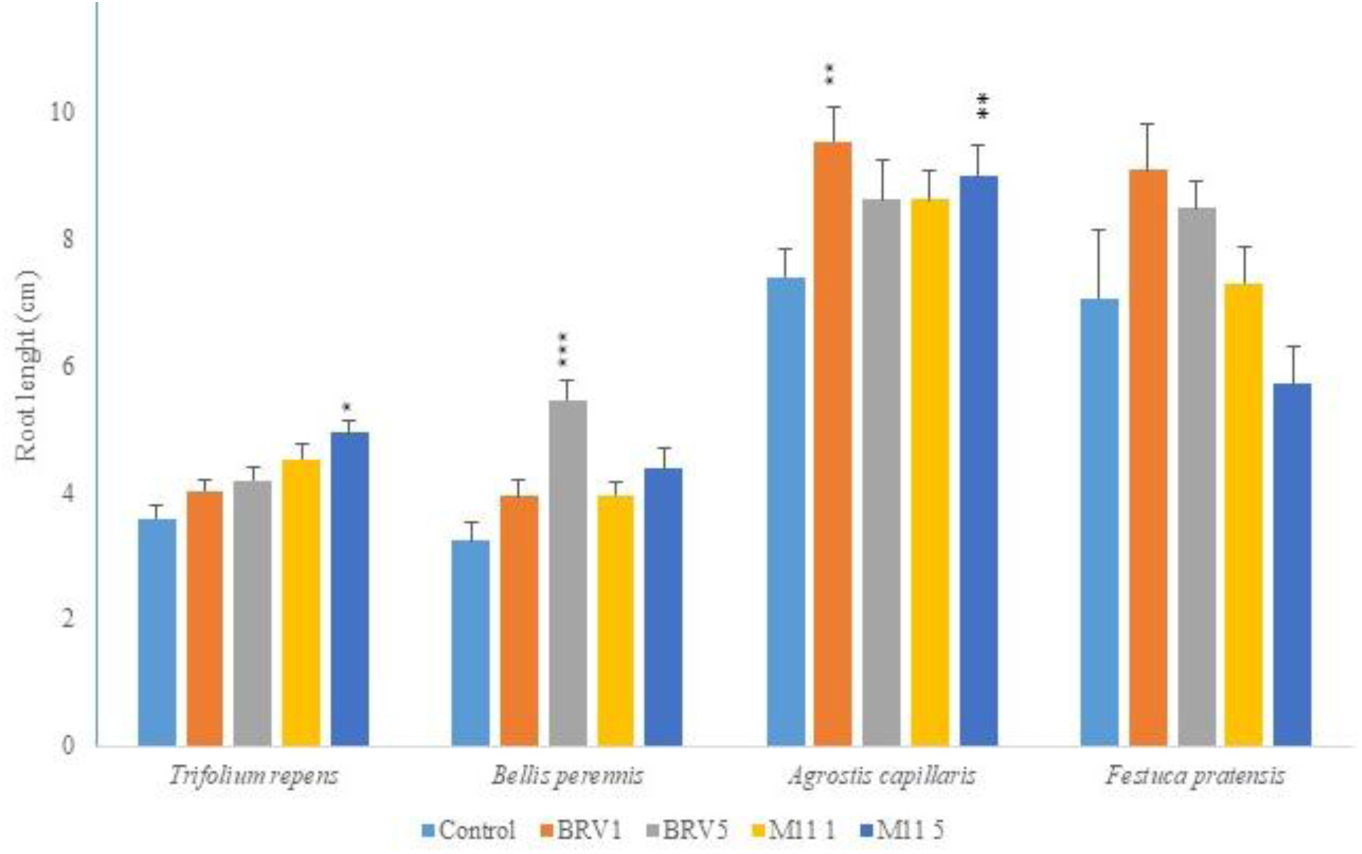
Effect of aqueous extracts on root length. Measures of root length of the different species in the different culture media. Data are presented as arithmetic means with their standard errors.

## 4. Discussion

This study examined the presence and biological activity of *Pteridium aquilinum* in northern Spain. Illudane glycosides were quantified in plant tissues and water samples from multiple bracken-infested sites, and the genotoxic and allelopathic effects of aqueous extracts were assessed. The detection of high concentrations of ptaquiloside in croziers from sites with reported livestock mortality underscores the real-world relevance of these findings, while the observed genotoxicity in Drosophila and human cell assays confirms the biological activity of plant-derived compounds. Interestingly, allelopathic assays indicated a potential stimulatory effect on the germination of meadow species, suggesting that bracken’s influence on plant communities may be more complex than simple inhibition. Collectively, these results highlight the need for targeted management strategies to mitigate both the ecological and health-related impacts of bracken proliferation.

The fern life cycle, comprising distinct gametophytic and sporophytic phases, is notoriously difficult to observe under natural conditions, especially in temperate regions where sporulation is rare. In northern Spain, bracken rarely produces spores, which limits opportunities to study early developmental stages in the field. In this context, in *vitro* culture provides a valuable approach to examine both phases separately and under controlled conditions (Sheffield et al., 2001; Eslava-Silva et al., 2009; Willyams and Daws 2014; Jang et al., 2020). In the present study, spores collected from Honduran plants germinated successfully *in vitro*, giving rise to gametophytes that produced both antheridia and archegonia, followed by sporophyte formation under appropriate nutritional conditions. Like previous reports in bracken and other fern species, nutrient strength and the ionic balance of the culture medium significantly influenced developmental outcomes: gametophytes proliferated rapidly on full-strength MS medium but rarely formed sporophytes, whereas half-strength MS medium promoted abundant sporophyte development (Kang et al., 2021). Conversely, sporophyte development in terms of frond number was stimulated by full-strength MS medium, although this was not the case for sporophyte length, which showed no improvement under full-strength conditions.

The detection of all four analyzed compounds: ptaquiloside (PTA), pterosin B (PtrB), pterosin A (PtrA), and ptesculentoside (PTE), in all crozier samples highlights the widespread occurrence of illudane glycosides in *P. aquilinum* populations from northern Spain. This observation is consistent with previous European studies reporting a broad geographical distribution of related glycosides in bracken ferns. For instance, Kisielius et al. (2024) analyzed PTA, CAU, and PTE across 66 sites in Denmark, Sweden, and Finland, identifying PTA as the predominant compound, with CAU and PTE occurring at lower relative abundances.

Earlier work by Sierra et al. (2025) demonstrated that pterosins A and B were present in all fourteen sampled Asturian sites, although with pronounced quantitative variability. In that study, PtrA levels ranged from 6 to 260 μg/g DW, while PtrB concentrations were approximately an order of magnitude higher. Extending this analysis to three additional sites, two of them associated with cattle mortality events (BRV and PEBEL), the present study shows that PtrA and PtrB concentrations remain within the ranges previously reported. As consistently observed, PtrB predominated over PtrA, reinforcing the notion that this pattern is conserved across different locations.

The finding of PtrA in plant material from all seventeen analyzed sites supports the widespread occurrence of its corresponding glycoside precursor, CAU, in Asturian bracken populations. Moreover, the identification of PTE in BRV, PEBEL, and MU2 is noteworthy, as this compound has historically been reported only sporadically in European bracken. Its presence in multiple sites suggests that PTE may be more common in northern Spanish populations than previously recognized. This is consistent with quantitative LC–MS studies showing that CAU and PTE can reach concentrations comparable to PTA in certain bracken populations (Kisielius et al., 2022), and with evidence indicating that both compounds may contribute to adverse effects in cattle (Fletcher et al., 2011; de Oliveira et al., 2020).

When restricting the analysis to the three newly studied sites, PTA levels, precursor of PtrB, were markedly higher at BRV compared to PEBEL and MU2, indicating a strong site-dependent variability. Although these concentrations did not reach the extreme values reported elsewhere (up to 45 mg/g DW; Kisielius et al., 2024), BRV consistently exhibited the highest levels of all four analyzed compounds. Together, the elevated concentrations of PTA, PtrB, PtrA, and PTE at this site may help explain the reported cases of cattle mortality associated with bracken consumption, in agreement with previous reports from Europe and other regions (Aranha et al., 2014).

Finally, the detection of CAU and PTE in both gametophyte and sporophyte generations derived from *in vitro*–cultured Honduran specimens indicates that the biosynthesis of these illudane glycosides is conserved across geographic origins and developmental stages. Furthermore, the elevated levels of PTA and PTE observed in *in vitro*–grown sporophytes, respect to the gametophytes, together with the detection of PTE in the PEBEL population from the Picos de Europa, suggest that both developmental stage and environmental or cultivation conditions play an important role in determining glycoside accumulation. This interpretation aligns with previous findings showing substantial variation in illudane glycoside content across life stages and growth conditions (Kisielius et al., 2022).

The ability of PTA to leach into both surface and groundwater raises concerns about environmental exposure and potential risks to livestock and humans. In this context, Rasmussen et al. (2016) investigated PTA in a stream draining a bracken-dominated catchment in Denmark, reporting base-flow concentrations below 61 ng/L, but storm-event peaks reaching up to 2.2 µg/L, demonstrating that rainfall can trigger pronounced pulses of PTA into surface waters. Complementarily, Skrbic et al. (2021) detected PTA in shallow groundwater wells within bracken-dominated areas at concentrations of approximately 0.35 µg/L. Expanding this perspective, Kisielius et al. (2022) examined the occurrence and stability of PTE, CAU, and PTA in surface waters across Denmark, Sweden, and Spain. In Asturias, PTE concentrations ranged from 0,4 to 5,3 µg/L and CAU from 0,3 to 2,3 µg/L, whereas PTA was not detected in surface-water samples.

In contrast, the results of the present study demonstrate the presence of PTA, PTE, and the pterosins PtrA and PtrB in water samples collected from ten bracken-infested areas across Asturias, indicating a broad geographical distribution. These findings highlight the environmental mobility of bracken-derived toxins and underscore the importance of monitoring both surface and groundwater in bracken-dominated regions. Given the broad spatial coverage of the sampling area and the analytical challenges associated with detecting these compounds in water, it is particularly noteworthy that all four metabolites were detected in half of the sampled sites, with at least two or three compounds detected in the remaining locations.

Genotoxicity analysis, *in vivo* in *Drosophila* e *in vitro* in cultured human cells, were performed in several of the collected samples. *In vivo* analysis with the water samples detected genotoxic activity when the samples were concentrated at least 3 or 4 times, especially in males, what might be relevant for animals or humans drinking it, depending on the consumed water volume or, independently of the volume, if some metabolites bioaccumulate in some tissues or organs of exposed individuals, such as it happens with metals, nanomaterials and some chemicals (Hou *et al*., 2016; Kershaw & Hall, 2019; Bhagat *et al*., 2020), or if water content might be concentrated at specific times along the year. When checking the results, the genotoxic activity was linked to the induction of somatic mutations, because mitotic recombination was not detected in any case. This finding did not agree with the genotoxic activity detected in aqueous extracts from Pteridium plants, in which recombination induction, although lower than mutation induction, was always detected (Sierra et al., 2025). Moreover, recombination induction is always detected on the SMART assay (Vogel and Nivard, 1993; 1999). Therefore, the lack of induced recombination with these samples should indicate the presence of some agents not present on the plants, that might interact with the plant lixiviates, or of some toxic plant agents (Gil da Costa et al., 2012-rev; Aranha et al., 2019; Malik et al., 2023), whose levels might accumulate in water, and interact with genotoxicity.

When the genotoxic activities were compared to the levels of detected metabolites no relationship was found, what means that the genotoxic activities of the samples were not dependent on the metabolite levels, and this agreed with the presence of other agents that might interact with genotoxicity. However, statistically significant relationships were found between the genotoxic activities of water samples and the genotoxic activities of the plants infesting the water sources, both in females and males. Therefore, since the genotoxic activities in plants were dependent on the metabolite levels, the lack of relationships between metabolites and genotoxic activities in water samples might be due to something lost, or degraded, on the water when the metabolites were determined (Kisielius et al., 2022).

The *in vivo* genotoxic analysis of the BRV plant sample revealed a genotoxic activity slightly higher than expected considering the altitude and latitude where it was collected, but lower than expected considering the levels of detected metabolites. Perhaps because of these unexpected results, when these data are combined with those previously published with other samples from Asturias, the described significant relationship with latitude (Sierra et al., 2025) was no longer found. However, statistically significant relationships between the genotoxic activity and the PtrA and PtrB contents were still detected, indicating that the contents of the illudane glycosides CAU and PTA, respectively, whether detected or not, were at least partially responsible for the detected genotoxicity.

In addition, since the *in vivo* genotoxic activity was related to metabolite content, it might be used to compare the risk associated with plants at different development stages, like young or senescent fronds, or to study the effect of the time used to prepare aqueous extracts. The obtained results showed a significantly lower genotoxic activity of senescent fronds, compared to young ones, suggesting a different metabolite content, or the degradation of metabolites (Kisielius et al., 2022). Moreover, when the plant was mixed with water to generate aqueous extracts, the time of this mix preparation was important for the extract genotoxic activity, and since it was different for females and males, it might influence recombination (Vogel and Nivard, 1999).

Genotoxic activity was also studied *in vitro* in cultured human cells. Results confirmed that aqueous plant extracts could generate DNA strand breaks, detected in the comet assay, as described before (Siman et al., 2000; Campos-da-Paz et al., 2008; Pereira et al., 2009). This damage was probably generated by alkylation of nitrogen atoms (Kushida et al., 1994; Freitas et al., 2001), through the generation of abasic sites that, in alkaline conditions, generated DNA single strand breaks (Beranek 1990; Kushida et al., 1994; O’Connor et al., 2019).

In view of the results on allelopathy, it seems that bracken extracts did not show any effect on the germination of the four analyzed species. Similar results have been reported in the literature (Bracho & Arnaud, 2012). After one month, seedlings were extracted from Petri dishes to collect data on root length and the number of leaves per seedling. While there were significant differences between treatments, there was no generalized pattern in these differences. It would have been expected that, due to a supposed allelopathic effect, root length and leaf number in the extracts would be lower than in the control. However, in cases where there were significant differences, the length and number of leaves in the extracts were always greater than in the control. A phytostimulant effect on some parameters has also been observed in an experiment conducted by Loresco et al. (2006), where *T. repens* seeds and perennial ryegrass were germinated with bracken extract. While germination was not affected, shoot growth was stimulated in all treatments compared to the control.

Stress-generating factors, such as drought, soil mineral deficiencies, low temperatures, or treatment with herbicides, could induce the fern to produce allelopathic compounds (Witt, 1999; Narwal, 1999). This could explain why no allelopathic effects were observed in the species tested; perhaps the samples used to make the extract did not develop allelopathic compounds because they were not exposed to stress. Another possible explanation is that during the extraction process, the compound(s) responsible for the allelopathic effect were not extracted, or were lost or degraded, leaving the extract as a nutrient-rich solution that may have favored the germination and development of the tested seedlings. Regarding the study of allelopathic effects, we cannot conclude that the extracts exert such an effect on the species studied. Although there was no statistical evidence, in some cases a slight tendency toward the promotion of root length and leaf number could be observed. Therefore, given these results under the tested conditions and species, other possible causes for the expansion of the common fern should be investigated beyond the potential allelopathic effects of its biochemical arsenal.

## 5. Conclusions

This study provides a comprehensive assessment of the distribution and biological activity of *Pteridium aquilinum*, quantifying the illudane glycosides PTA and PTE and their metabolic products, pterosins A and B, across *in vitro-*cultured gametophytes and sporophytes, spring-collected croziers, and autumnal water samples, thereby integrating plant tissues and environmental matrices. The use of laboratory standards enabled accurate compound quantification, while genotoxicity assays revealed the potential induction of mutations and recombination events in somatic cells of a higher eukaryotic organism *in vivo*. In addition, allelopathic bioassays suggested possible stimulatory effects on meadow species, pointing to a more complex ecological role of bracken than previously recognized. Despite its ecological and veterinary relevance, research on *P. aquilinum* in Spain remains limited and fragmented. Addressing this gap through systematic toxin monitoring, targeted epidemiological studies, and the development of sustainable management strategies is essential to safeguard cattle health, rural economies, and potentially human health, given the well-established carcinogenic potential of bracken-derived toxins.

## Acknowledgments

The authors thank Jos for his invaluable help with the logistics with sampling. We would like to extend our gratitude to The National Institute of Forest Conservation and Development, Protected Areas and Wildlife (ICF), Ministry of Natural Resources and Environment (SERNA), and Friends of La Tigra Foundation (AMITIGRA), as well as La Tigra National Park Management Plan (PNLT) 2013–2025. Tegucigalpa, Honduras.

## Author Contributions

Conceptualization, HF. and LMS.; Methodology, IF, LMS, AEP, MLR, BB, II, AV, AB, DG, JMA.; Formal analysis, IF, LMS, Statistical analysis, LMS, and JMA, Writing—original draft, HF and LMS, Writing—review and editing, LMS, HF, IF, MLR, AEP and JMA; Project administration, HF, Funding acquisition, HF, LMS, IF, JM, EC and JMA. All authors have read and agreed to the published version of the manuscript.

## Funding

This research was funded by the Spanish Ministry of Science and Innovation, Reference TED131270B-100, in the Recovery, Transformation and Resilience Plan.

## Notes

### Competing Interest Statement

The authors have declared no competing interest.

