## supplemental material for "Unveiling the Toxicological and Allelopathic Effects of *Pteridium aquilinum*: Chemical Profiling and Biological Assays"

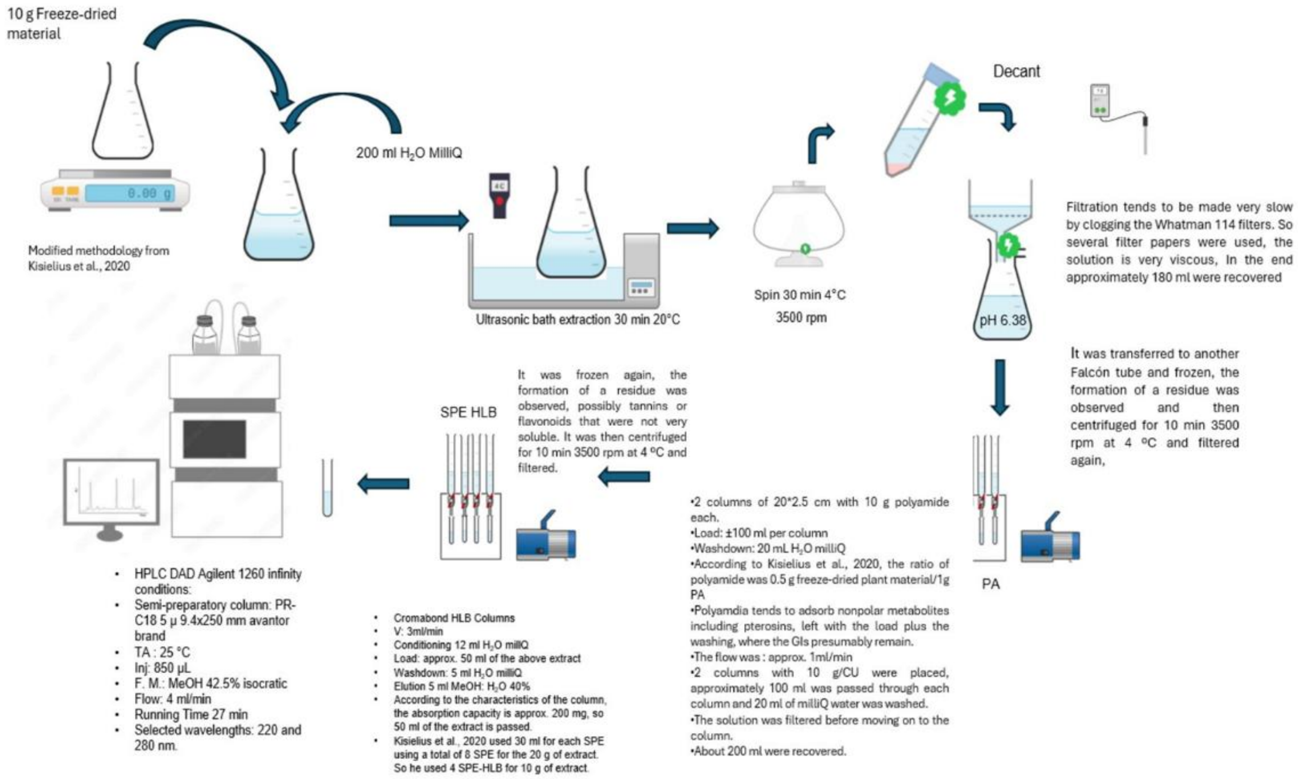


**Figure S1.** Scheme of the isolation and purification of illudane glycosides from *Pteridium sculentum*.

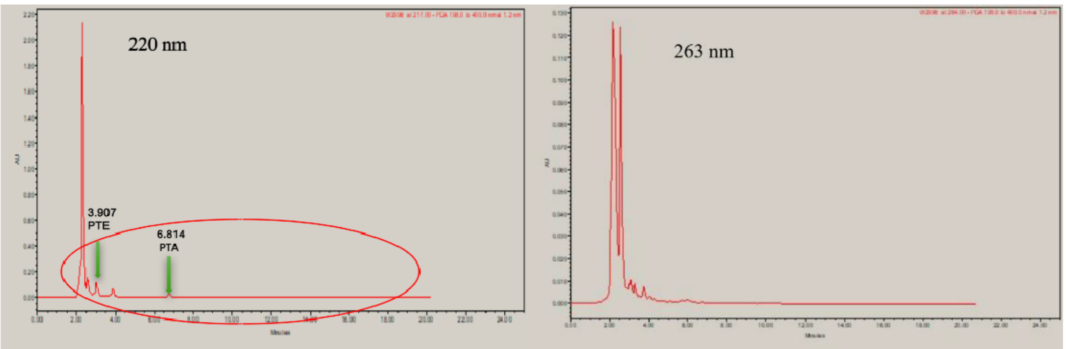


**Figure S2.** Chromatograms obtained from the aqueous extract of *Pteridium esculentum* passed through polyamide resin at 220 nm and 263 nm.

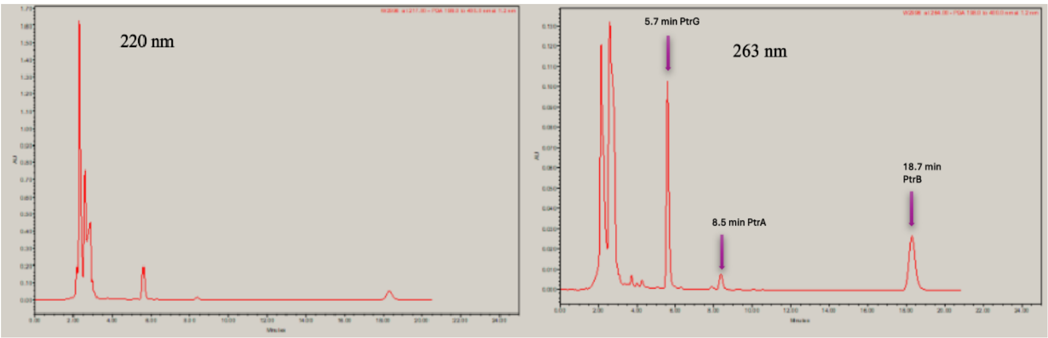


**Figure S3.** Chromatograms obtained from the aqueous extract of *Pteridium esculentum* passed by polyamide and hydrolyzed at 220 nm and 263 nm. The signals that are lost are identified and the signals corresponding to the pterosins are observed again.

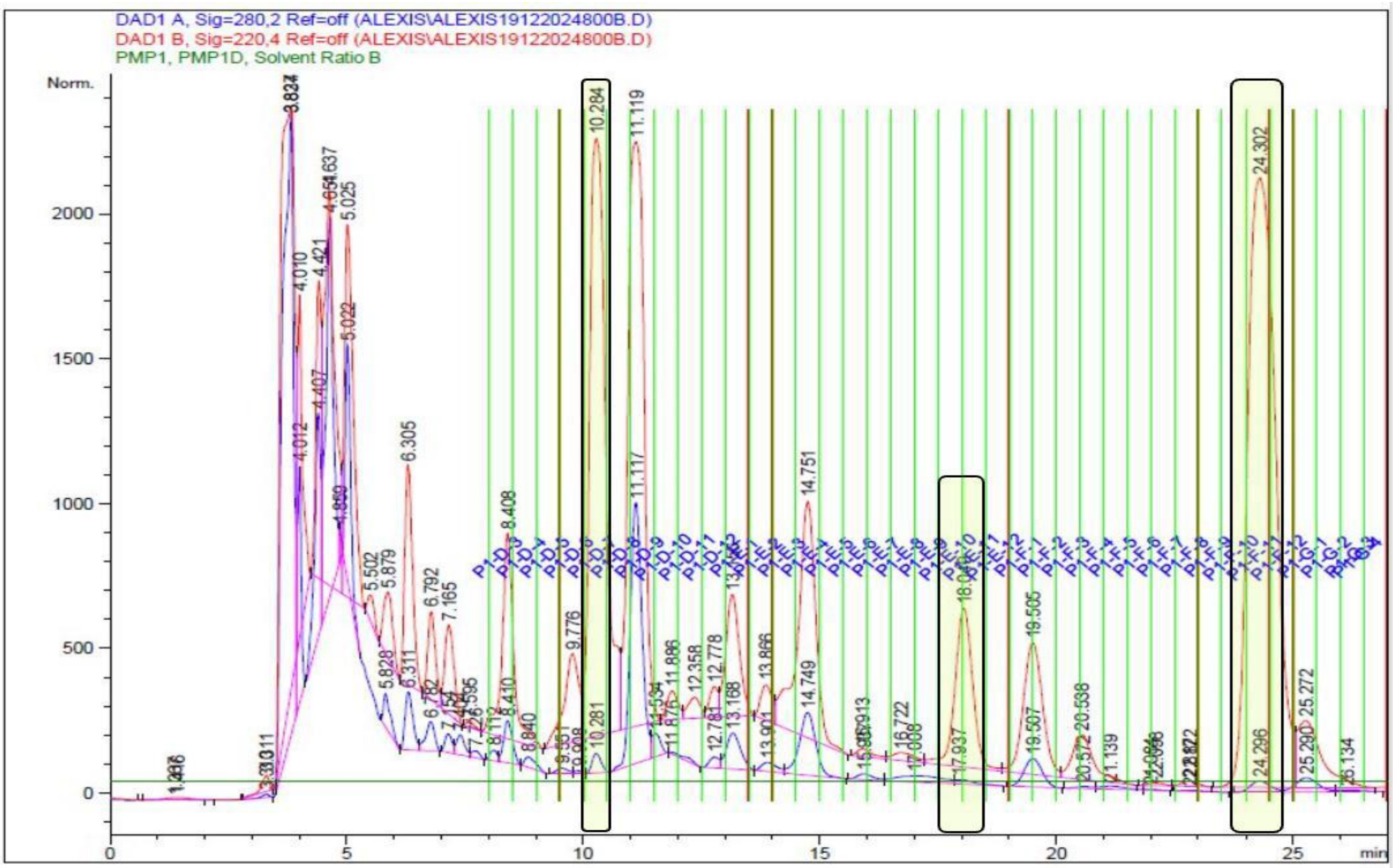


**Figure S4.** Fractional chromatogram of the equipment, the highlighted fractions correspond to the absorption peaks reported in the literature.

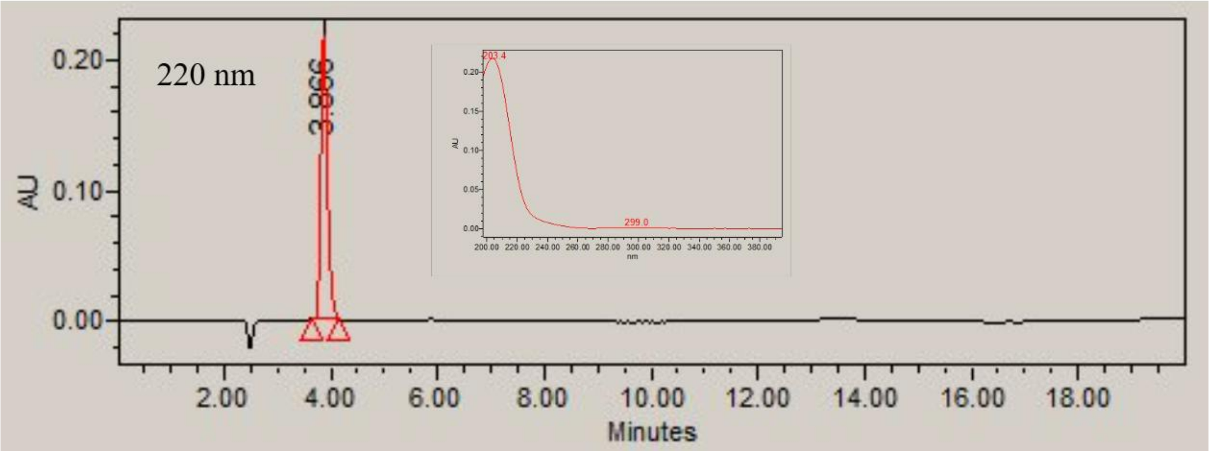


**Figure S5.** Chromatogram of the **R01 (PTE)** fraction with HPLC-PAD and absorption spectrum.

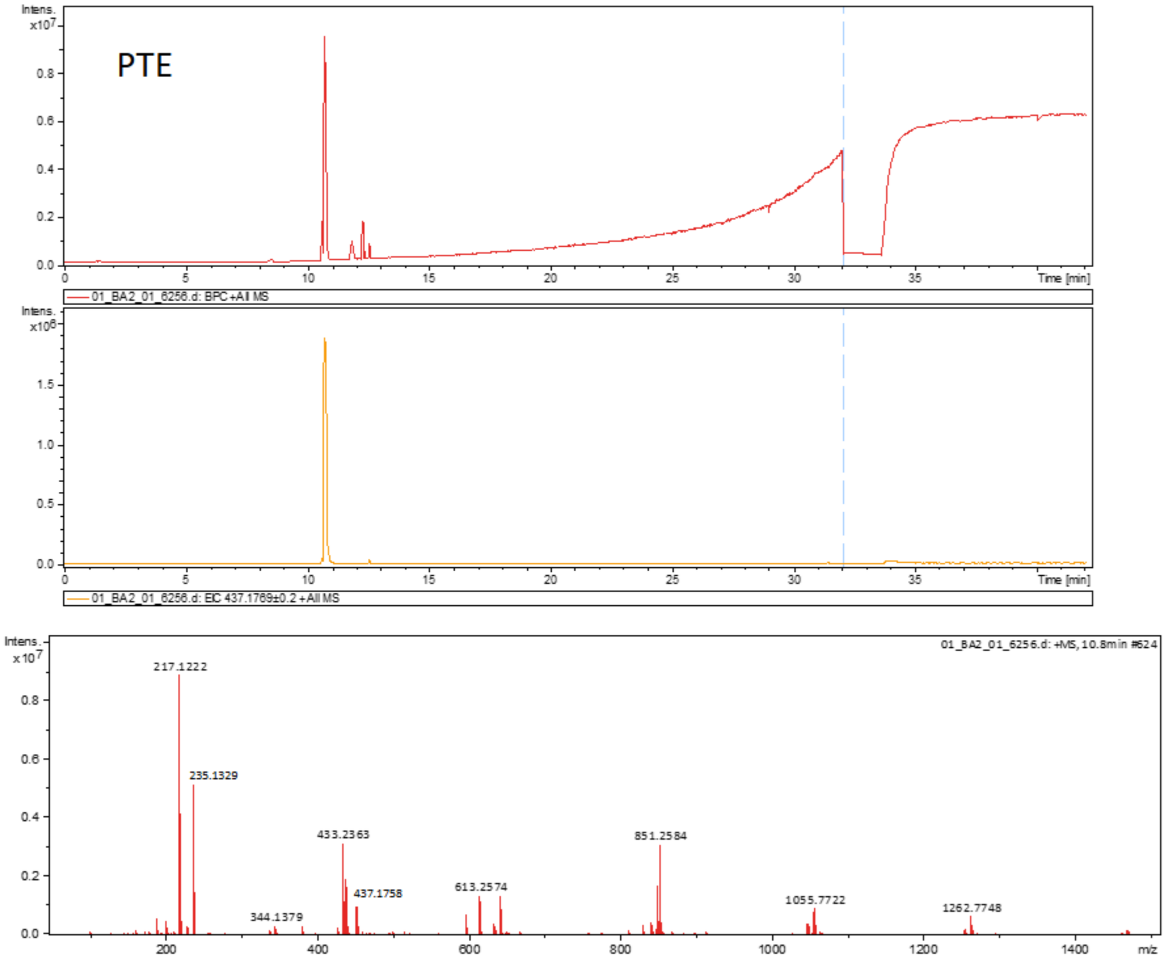


**Figure S6.** Chromatogram of the **R01 (PTE)** sample performed with UPLC-HRMS.

### High Resolution Mass Spectrum

**Analysis Info**

| | | | | |
|---|---|---|---|---|
| Method | MS_MASA_EXACTA.m | Operator | Operator | |
| Sample Name | FRACC_R01 | Instrument | impact II | 1825265.10101 |
| Comment | | | | |

**Acquisition Parameter**

| | | | | | |
|---|---|---|---|---|---|
| Source Type | ESI | Ion Polarity | Positive | Set Nebulizer | 2.4 Bar |
| Focus | Active | Set Capillary | 4000 V | Set Dry Heater | 250 °C |
| Scan Begin | 50 m/z | Set End Plate Offset | -500 V | Set Dry Gas | 6.0 l/min |
| Scan End | 1500 m/z | Set Charging Voltage | 2000 V | Set Divert Valve | Source |
| | | Set Corona | 0 nA | Set APCI Heater | 0 °C |

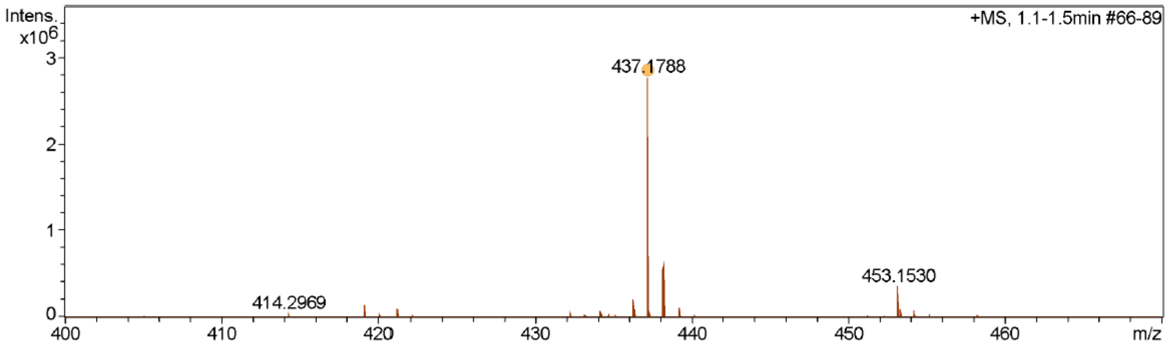


| Meas. m/z | # | Ion Formula | m/z | err [ppm] | mSigma | # mSigma | Score | rdb | e⁻ Conf | N-Rule | Adduct |
|---|---|---|---|---|---|---|---|---|---|---|---|
| 437.1788 | 1 | C20H30NaO9 | 437.1782 | -1.5 | 3.1 | 1 | 100.00 | 6.0 | even | ok | M+Na |

**Figure S7.** High-resolution mass analysis of the **R01 (PTE)** sample.

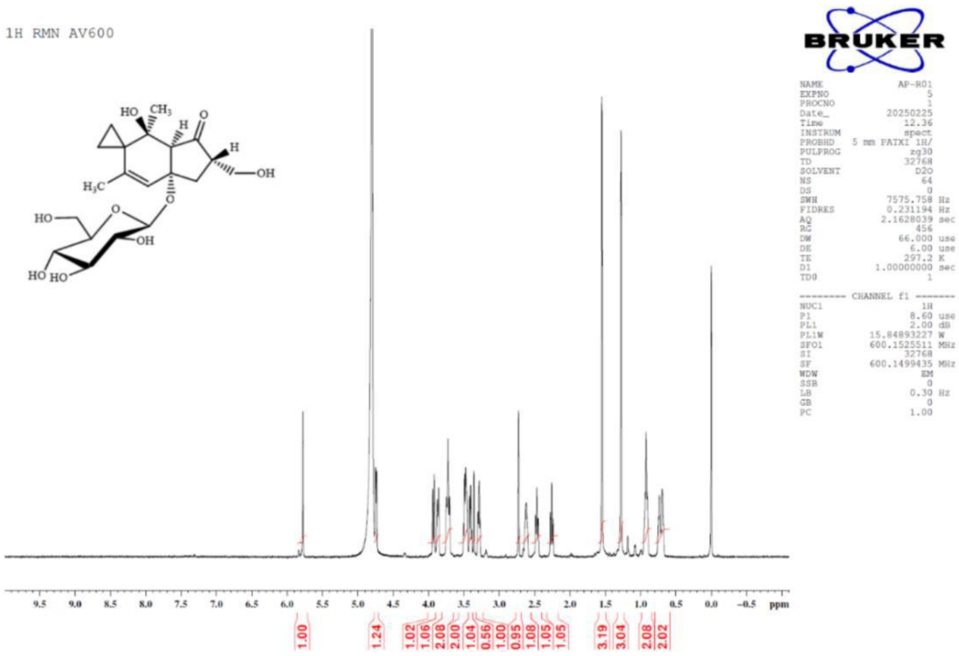


**Figure S8.** NMR-$^{1}$H spectrum of the R01 (PTE) sample.

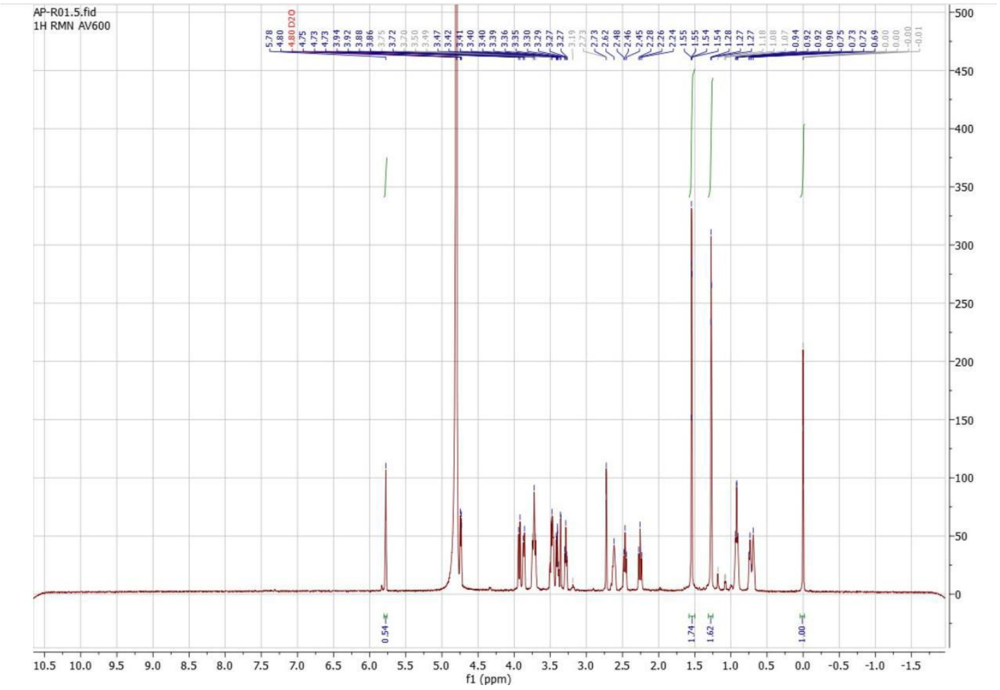


**Figure S9.** NMR-$^{1}$H spectrum showing the relative integral of allyl and methyl hydrogens.

**The purity.**

The purity of the analyte was determined using the following equation:

$$M_x = \frac{Al_x}{Al_i} \cdot \frac{NN_i}{NN_x} \cdot n_i \cdot MW_x \cdot P_i$$

where ***x*** and ***i*** denote the analyte and the internal standard, respectively; ***Al*** represents the relative integral of the selected NMR signals; ***NN*** corresponds to the number of nuclei; n is the number of moles; ***MW*** is the molecular weight (g/mol); and $\boldsymbol{P_i}$ is the purity of the internal standard. The percentage purity of the analyte was subsequently calculated by comparing the calculated mass to the weighed sample mass:

$$\%\text{Purity}_x = \frac{M_x}{m_{\text{sample}}} \cdot 100$$

where $\boldsymbol{M_x}$ is the mass obtained from the first equation and $\boldsymbol{m_{samplem}}$ is the mass of the sample used for the proton NMR analysis.

**Table S1** Purity of the **R01 (PTE)** sample.

| **Sample Mass:** 1.10 mg | **Proton used** | % Purity of **PTE** |
|---|---|---|
| ***AI$_x$/AI$_i$**: 1.74/1 | H (δ 1.55 ppm) | 86.00% |
| ***AI$_x$/AI$_i$**: 0.54/1 | H (δ 5.78 ppm) | 80.20% |
| ****Purity of the Internal Standard** 99.9% $\boldsymbol{n_i}$ 4.38 x $10^{-4}$ *mmols* | **Average** | 83.10% |

* Data obtained from the $^{1}$H-NMR spectrum.

** 3-(trimethylsilyl)propionic-2,2,3,3-d4 acid, sodium salt (**internal standard**) used in $^{1}$H NMR.

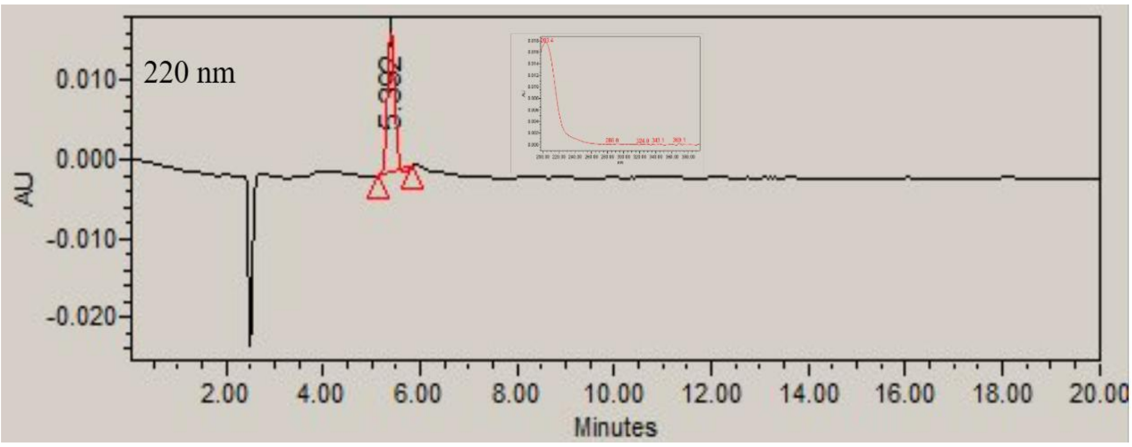


**Figure S10.** Chromatogram of the **R02 (CAU)** sample in the HPLC-PAD and absorption spectrum.

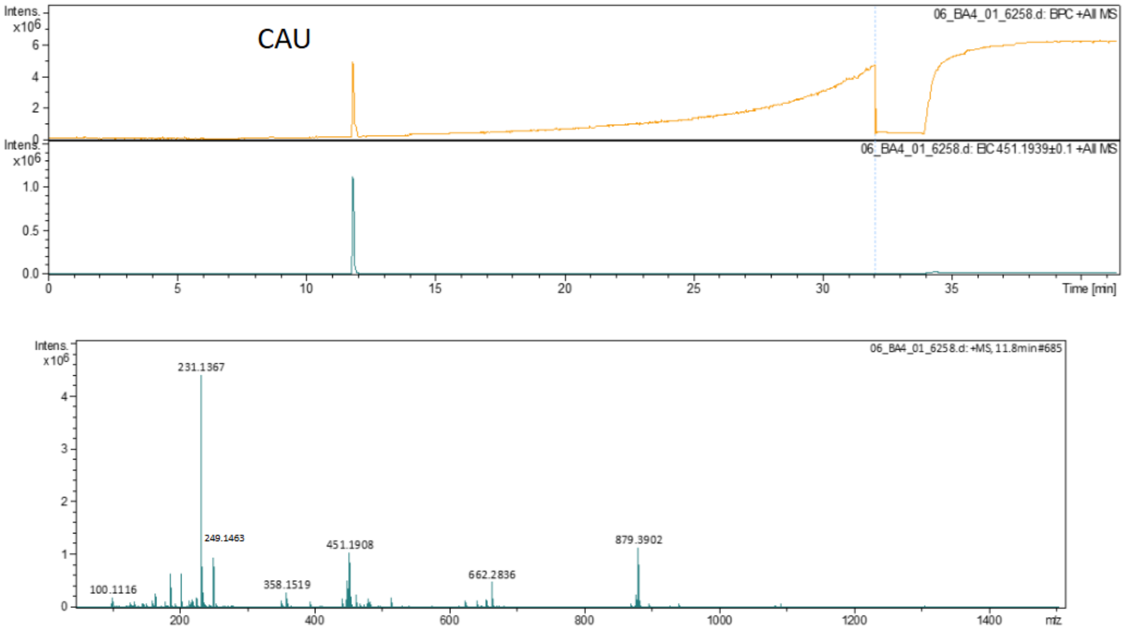


**Figure S11.** Chromatogram of the **R02 (CAU)** sample performed using UPLC-HRMS.

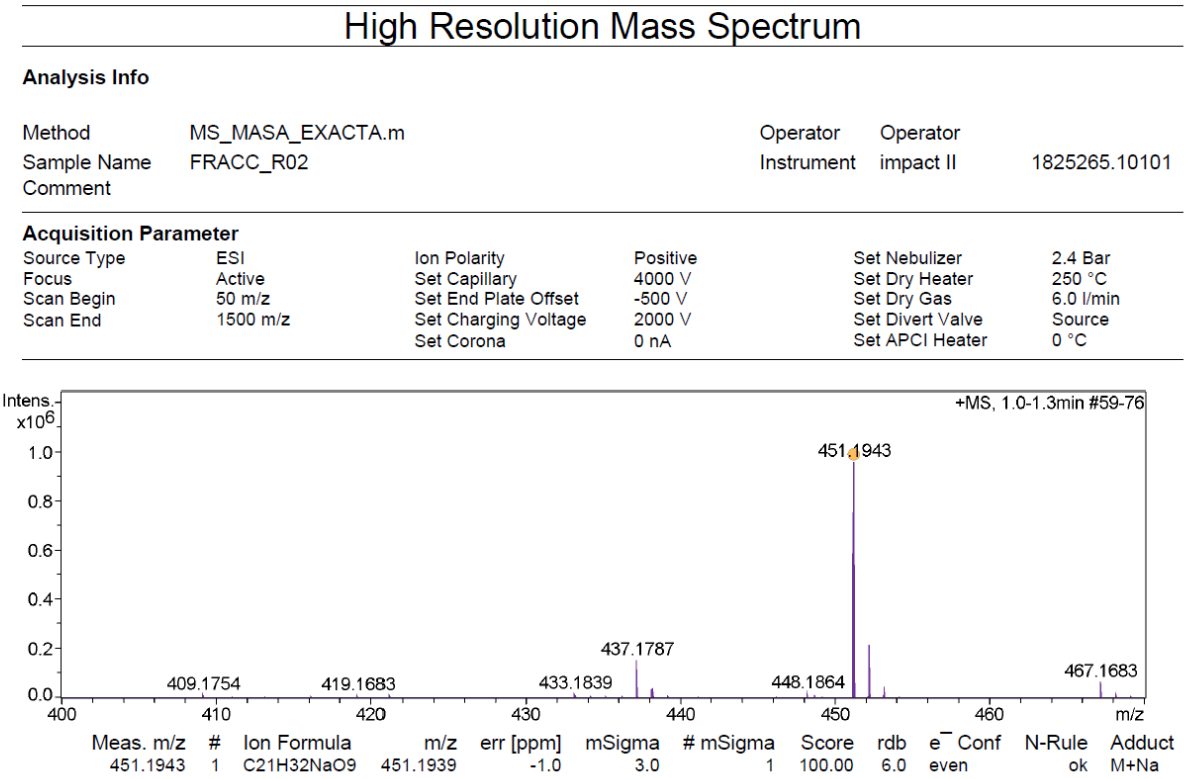


**Figure S12.** High-resolution mass analysis of the **R02 (CAU)** sample.

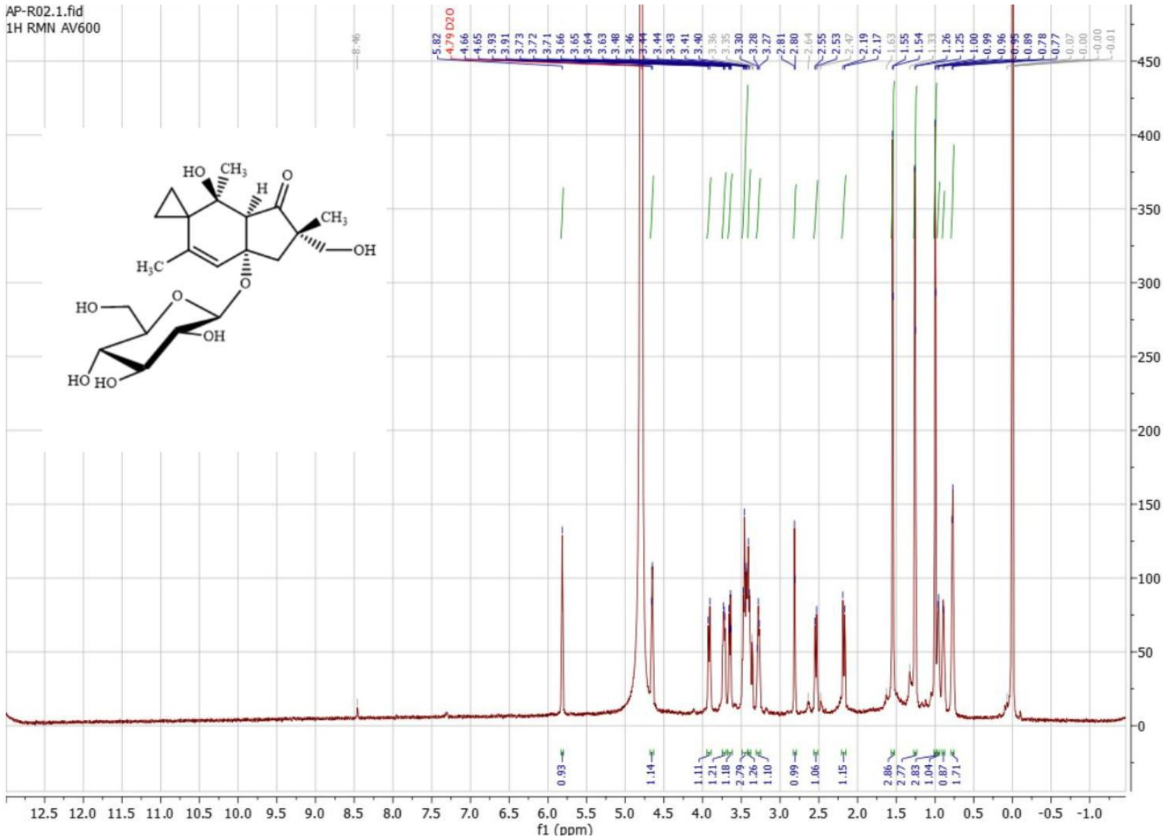


**Figure S13.** NMR-$^{1}$H spectrum of the **R02 (CAU)** sample.

**Table S2.** Purity of the **R02 (CAU)** sample.

| **Sample Mass:** 0.50 mg | **Proton used** | **% Purity of the CAU** |
|---|---|---|
| ***$AI_x/AI_i$**: 0.67/1 | H (δ 1.54 ppm) | 75.30% |
| ***$AI_x/AI_i$**: 0.17/1 | H (δ 5.76 ppm) | 57.30% |
| ****Purity of the Internal Standard** 99.9% $n_i$ 4.38 x $10^{-4}$ *mmols* | **Average** | 66.32% |

* Data obtained from the $^1$H-NMR spectrum.

** 3-(trimethylsilyl)propionic-2,2,3,3-d4 acid, sodium salt (**internal standard**) used in $^1$H NMR.

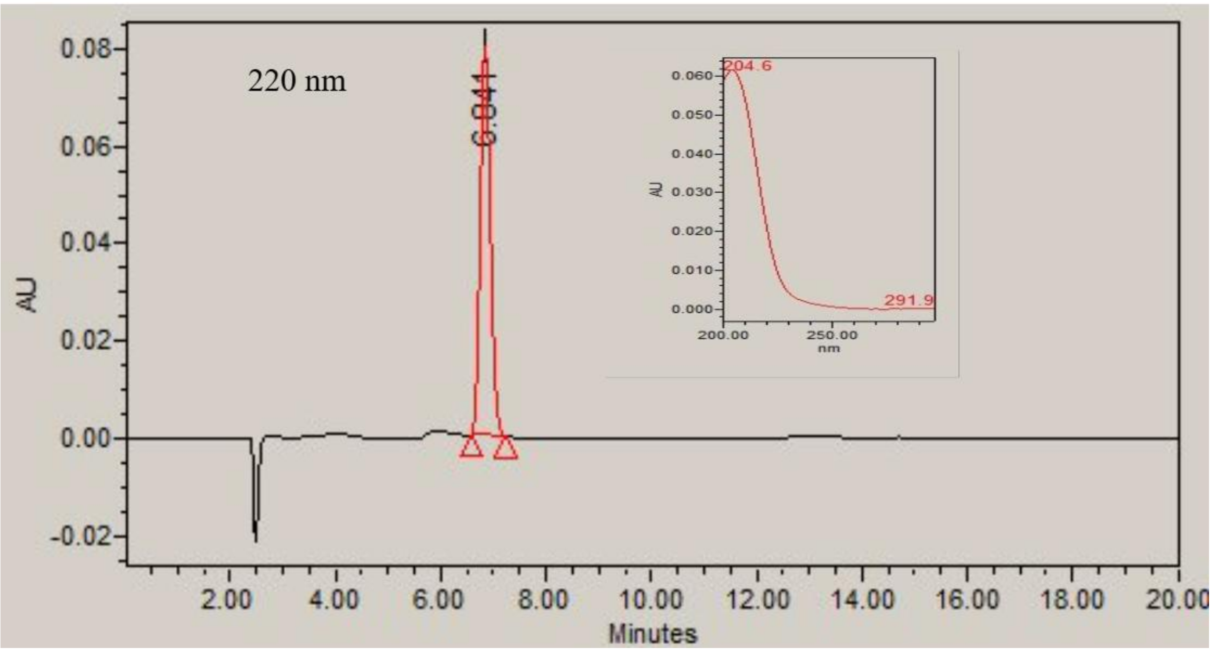


**Figure S14**. Chromatogram of the **R03 (PTA)** sample performed by HPLC-PAD at 204 nm and absorption spectrum.

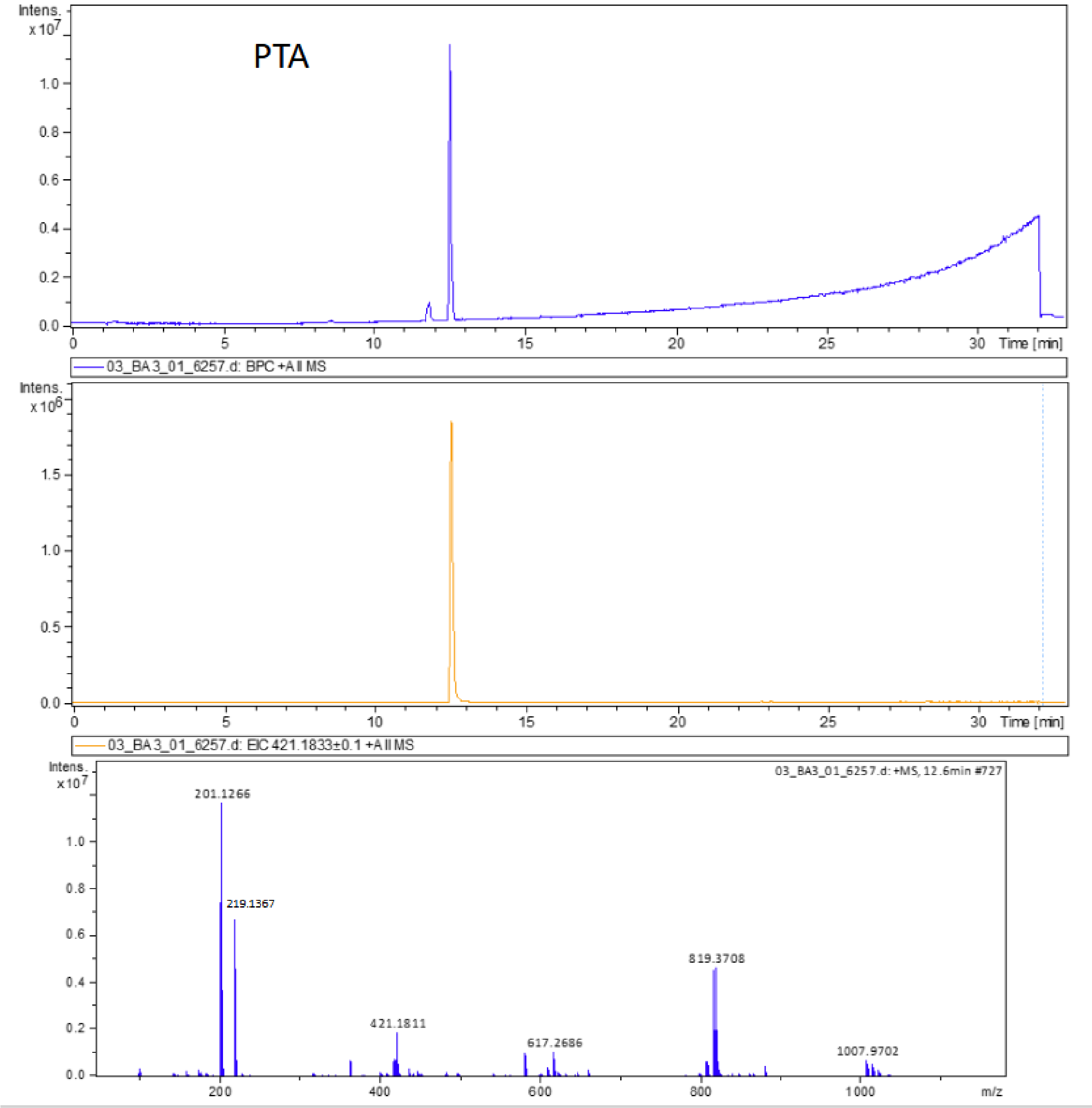


**Figure S15.** Chromatogram of the R03 (PTA) sample was performed using UPLC-HRMS.

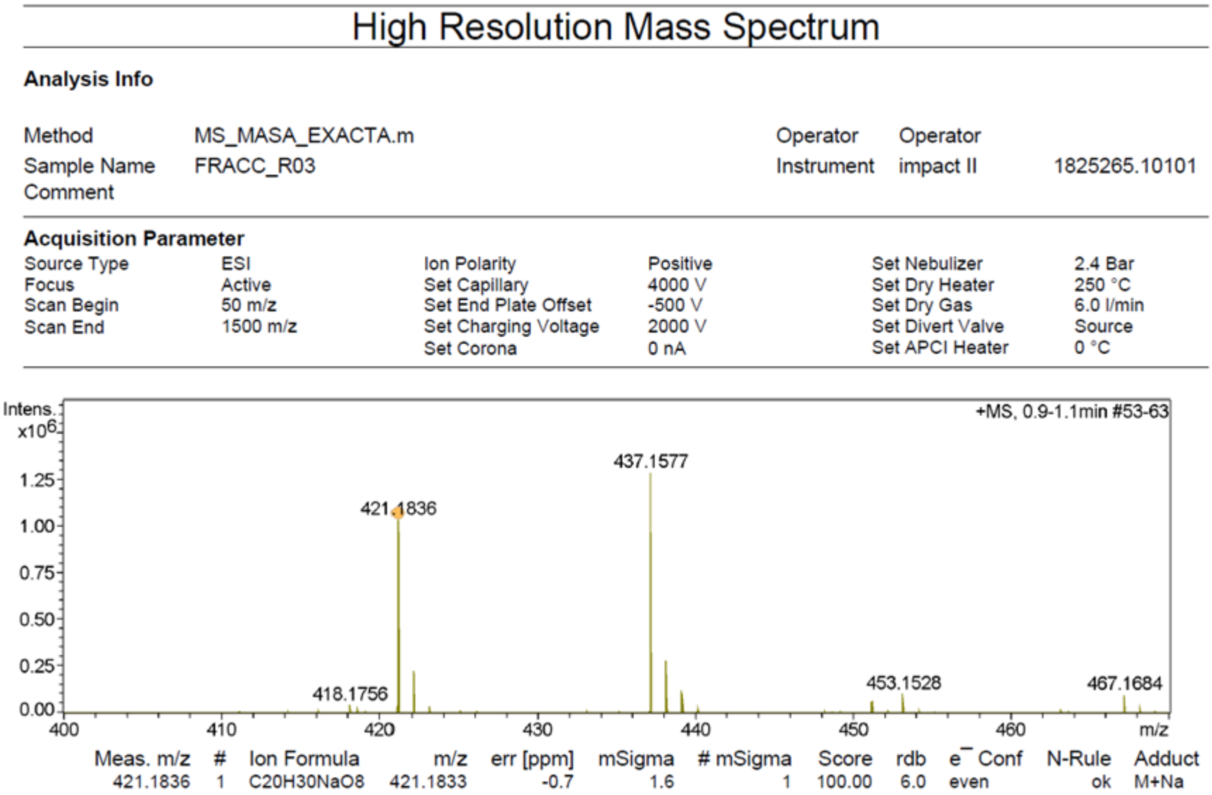


**Figure S16**. High-resolution mass analysis of the **R03 (PTA)** sample.

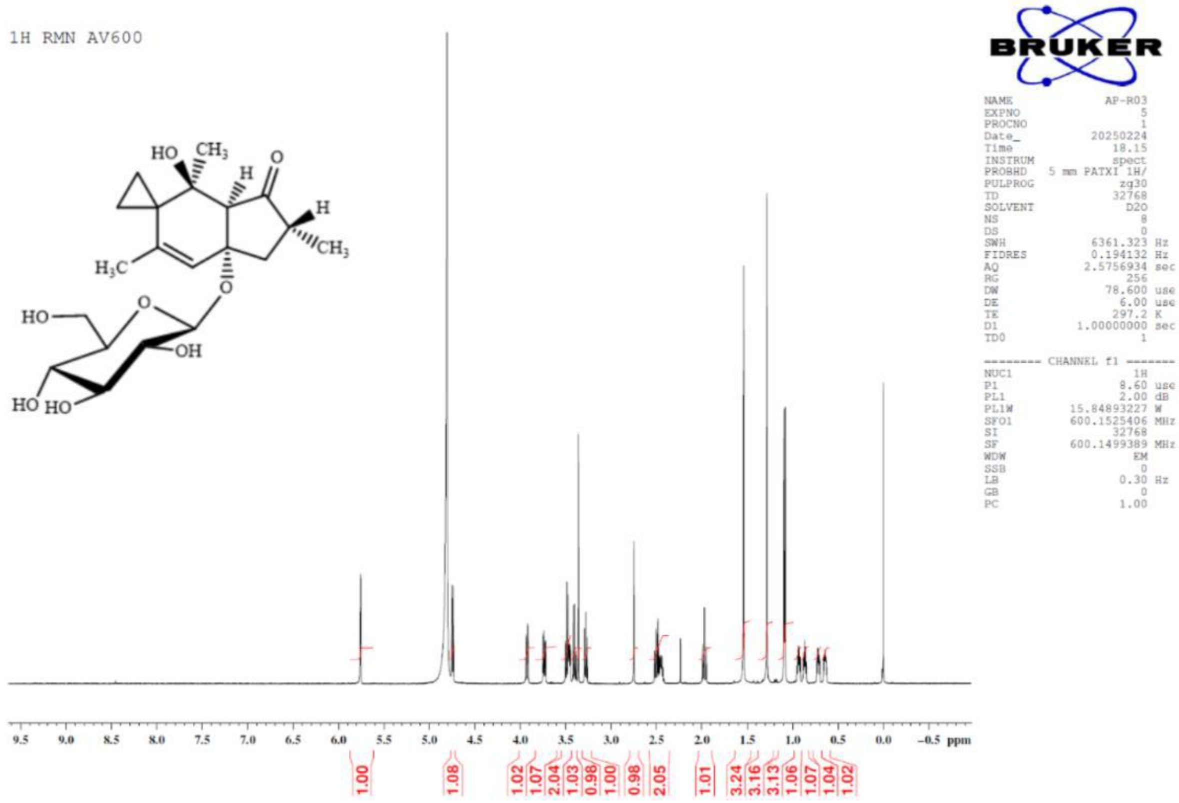


**Figure S17.** NMR-$^{1}$H spectrum of the **R03 (PTA)** sample.

**Table S3.** Purity of the **R03 (PTA)** sample.

| **Sample Mass:** 0.50 mg | **Proton used** | **% Purity of PTA** |
|---|---|---|
| ***$AI_x/AI_i$**: 2.39/1 | H (δ 1.54 ppm) | 96.16% |
| ***$AI_x/AI_i$**: 0.75/1 | H (δ 5.76 ppm) | 90.50% |
| ****Purity of the Internal Standard** 99.9% $n_i$ 4.38 x $10^{-4}$ *mmols* | **Average** | 93.33% |

* Data obtained from the $^1$H-NMR spectrum.

** 3-(trimethylsilyl)propionic-2,2,3,3-d4 acid, sodium salt (**internal standard**) used in $^1$H NMR.

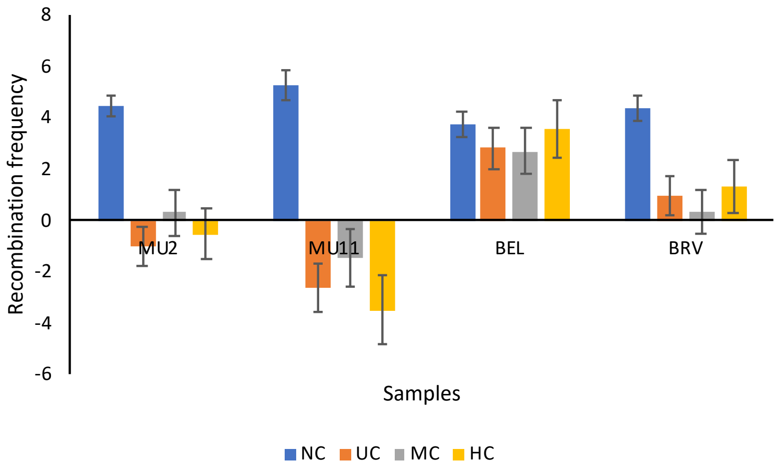


**Figure S18. Induction of recombination.** Induced recombination frequencies estimated subtracting the frequencies of mosaic eyes induced in males from those induced in females
